# Maximum likelihood estimation of species trees from gene trees in the presence of ancestral population structure

**DOI:** 10.1101/700161

**Authors:** Hillary Koch, Michael DeGiorgio

## Abstract

Though large multilocus genomic datasets have led to overall improvements in phylogenetic inference, they have posed the new challenge of addressing conflicting signals across the genome. In particular, ancestral population structure, which has been uncovered in a number of diverse species, can skew gene tree frequencies, thereby hindering the performance of species tree estimators. Here we develop a novel maximum likelihood method, termed TASTI, that can infer phylogenies under such scenarios, and find that it has increasing accuracy with increasing numbers of input gene trees, contrasting with the relatively poor performances of methods not tailored for ancestral structure. Moreover, we propose a supertree approach that allows TASTI to scale computationally with increasing numbers of input taxa. We use genetic simulations to assess TASTI’s performance in the four-taxon setting, and demonstrate the application of TASTI on a six-species Afrotropical mosquito dataset. Finally, we have implemented TASTI in an open-source software package for ease of use by the scientific community.

---

Large multilocus datasets are becoming ever more common in systematic biology (Cranston et al. 2009; Song et al. 2012; McCormack et al. 2013; Salichos and Rokas 2013; DeGiorgio et al. 2014; Fontaine et al. 2015; Peters et al. 2017). However, even with these more substantial datasets, the presence of gene tree discordance, in which inferred gene tree topologies conflict across loci, can be problematic when making phylogenetic inference. A major source of gene tree discordance is incomplete lineage sorting, which occurs when sampled lineages fail to coalesce, or find a common ancestor, in the first population in which they are able to do so. Accordingly, several statistically consistent methods for species tree inference that are robust to incomplete lineage sorting under the multispecies coalescent model have been developed (*e.g.*, Kubatko et al. 2009; Degnan et al. 2009; Liu et al. 2010b,c; Mossel and Roch 2010; Jewett and Rosenberg 2012; Helmkamp et al. 2012; Wu 2012; Mirarab et al. 2014; Mirarab and Warnow 2015).

The multispecies coalescent assumes that each modern and ancestral species is unstructured and has a constant population size, and that each pair of lineages within a given ancestral species has an equal probability of coalescing (Nakhleh 2013). Under these assumptions, incomplete lineage sorting leads to symmetries in gene tree distributions for any species tree, regardless of the number of taxa. For example, if a pair of taxa A and B are sister species on a species tree, then for all other species *X*, species A and *X* and species B and *X* have the same probability of being sister taxa in a gene tree (Allman et al. 2011). In the face of some form of gene flow between species, such as hybridization, continuous migration, or horizontal gene transfer, asymmetries in gene tree distributions can arise (Yu et al. 2011, 2012; Leache et al. 2014; Tian and Kubatko 2016; Long and Kubatko 2018). As such, asymmetries in gene tree distributions are often attributed to gene flow between species (McGuire et al. 2007; Escobar et al. 2012; Marcussen et al. 2014).

Despite the common crediting of asymmetries in gene tree distribution to inter-species gene flow, they can also emerge in the absence of such gene flow when ancestral populations are structured (Slatkin and Pollack 2008). There are several hypothesized examples of structured ancestral species (*e.g*., Garrigan et al. 2005; Thalmann et al. 2007; White et al. 2009), and genomic signatures of ancestral structure have been uncovered in a number of diverse lineages, including mouse (White et al. 2009) and yeast (Yu et al. 2012). Still, many methods for inferring species trees from multilocus data are not robust to ancestral structure, and can be proven to be positively misleading (DeGiorgio and Rosenberg 2016). DeGiorgio and Rosenberg (2016) demonstrate that exceptions to this rule are the maximum likelihood estimators of the species tree implemented in GLASS (Mossel and Roch 2010), STEM (Kubatko et al. 2009), and Maximum Tree (Liu et al. 2010c); however, these algorithms underperform on empirical data when gene trees are inferred rather than known.

Here we detail an algorithm named TASTI (Taxa with Ancestral structure Species Tree Inference), a maximum likelihood method for inferring species tree topologies when ancestral species are structured. We demonstrate that by explicitly incorporating the potential for population structure into a multi-species coalescent framework, the method is robust to population structure, and its performance typically does not significantly erode when applied to empirical data. While the method was designed to address incomplete lineage sorting caused by ancestral population structure, we demonstrate that it is also effective under the standard unstructured multispecies coalescent model. Finally, we propose an approach to infer species tree topologies under ancestral population structure for an arbitrarily large number of species. We show via simulations in the three- and four-taxon settings that TASTI outperforms competing methods MP-EST (Liu et al. 2010b), STELLS2 (Pei and Wu 2017), and STEM2.0 (Kubatko et al. 2009) when population structure is present, but remains competitive under the unstructured multi-species coalescent.

## Methods

### Modeling three species with ancestral population structure

In this section we describe a model relating three species with population structure in each ancestral population, and use it to construct probabilities of gene trees given a set of model parameters. In our model, the topologies ((*ab*)*c*) and ((AB)C) denote a gene tree and species tree, respectively, with sister species A and B and outgroup C. Figure 1a depicts a species tree with topology ((AB)C), in a configuration such that species B and C descend from the same ancestral subpopulation.

**Figure 1:**
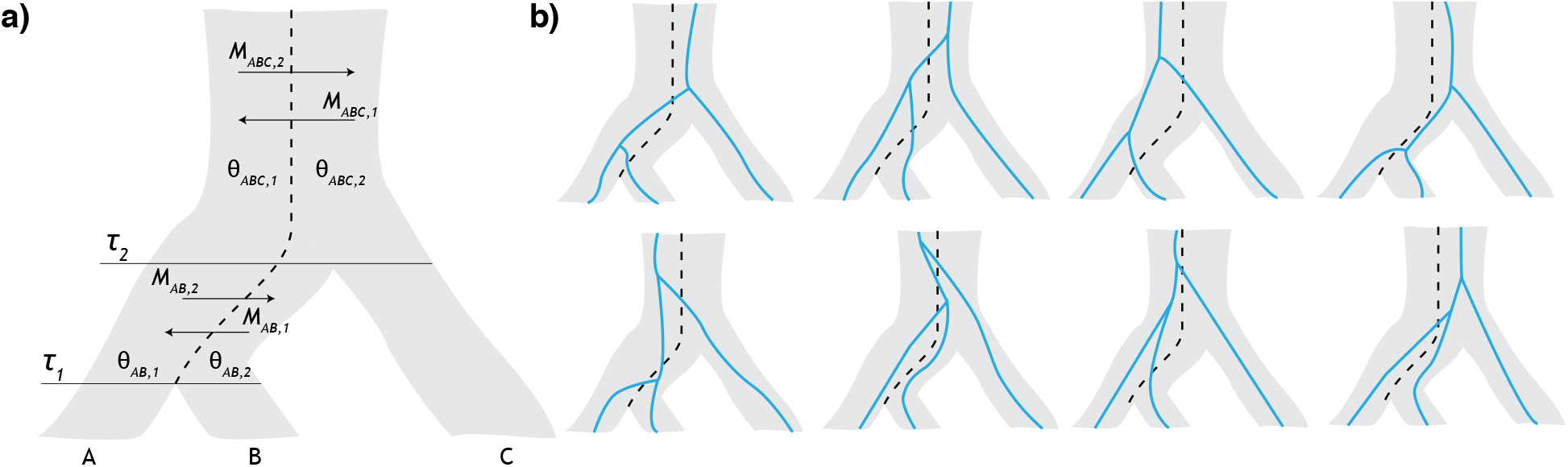
Modeling ancestral population structure. *a*) Model of the relationships among species A, B, and C with divergence times *τ*_1_ and *τ*_2_. Ancestral species belong to one of two subpopulations of scaled coalescent rates *θ*_*X,k*_ with migration between subpopulations at rates *M*_AB,1_ and *M*_AB,2_ below the root and *M*_ABC,1_ and *M*_ABC,2_ above the root. Lineages from species A descend from subpopulation 1, whereas lineages from species B and C descend from subpopulation 2. *b*) Incorporating population structure into the model increases the number of possible paths lineages from species A, B, and C might take to find their most recent common ancestor. For example, assuming that lineages from species A merge into subpopulation 1 and lineages from species B and C merge into subpopulation 2, a gene tree where lineages from species A and B coalesce first may arise from eight possible histories, as depicted here.

The species tree in our two-subpopulation model is characterized as follows. Divergence times, measured in expected number of mutations per site, are denoted ***τ*** = (*τ*_1_, *τ*_2_), where the internal branch length *τ*_2_ − *τ*_1_ is of particular interest. For all ancestral species *X*, *θ*_*X,j*_ = 4*N*_*X,j*_*μ*, where *N*_*X,j*_ is the effective population size of subpopulation *j* in ancestral species *X*, and *μ* is the mutation rate per site per generation. The mutation-scaled coalescent rates within subpopulations of ancestral species are inversely proportional to their respective values described in the vector ***θ*** = (*θ*_AB,1_, *θ*_AB,2_, *θ*_ABC,1_, *θ*_ABC,2_). Migration occurs between subpopulations at mutation-scaled rates according to the vector **M** = (*M*_AB,1_, *M*_AB,2_, *M*_ABC,1_, *M*_ABC,2_), where *M*_*X,j*_ is the rate at which lineages move to subpopulation *j* when in ancestral species *X*. Specifically, we define *M*_*X,j*_ = 4*N*_*X,j*_*m*_*X,j*_/*θ*_*X,j*_, where *m*_*X,j*_ is the fraction of subpopulation *j* in ancestral species *X* made up of migrants from its complementary subpopulation each generation.

Our method partitions the species tree into three distinct time transects on the intervals [0, *τ*_1_), [*τ*_1_, *τ*_2_), and [*τ*_2_, ∞). During the interval [0, *τ*_1_), we assume extant populations to be unstructured. Along the internal branch [*τ*_1_, *τ*_2_), we introduce ancestral subpopulations such that migration and coalescence events may occur between the two lineages from sister species A (originating in subpopulation 1) and B (originating in subpopulation 2). In the final interval [*τ*_2_, ∞), the lineage from species C is introduced into subpopulation 2. This setting matches the one presented by Slatkin and Pollack (2008), up to the first coalescence event. Migration and coalescence events occur until lineages from each species find their most recent common ancestor (MRCA), which may only happen between two lineages when they are in the same subpopulation. Since we are in essence estimating the waiting times until some occurrence, the distribution of migration and coalescent events can be modeled using an exponential distribution parametrized by an instantaneous rate matrix (Hobolth et al. 2011; Tian and Kubatko 2016). This parameterization yields the matrix exponential 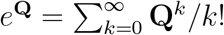, whose (*i, j*)th entry is denoted (*e*^**Q**^)_*ij*_.

### Instantaneous rate matrices for the model

The migration and coalescsence of lineages within a structured population can be explicitly modeled using instantaneous rate matrices. For time to the first coalescence *T*_1_ ∈ [*τ*_1_, *τ*_2_), where migration and coalescence can only occur for a pair of lineages, one from species A and one from species B, we define the instantaneous rate matrix **Q**_AB_.

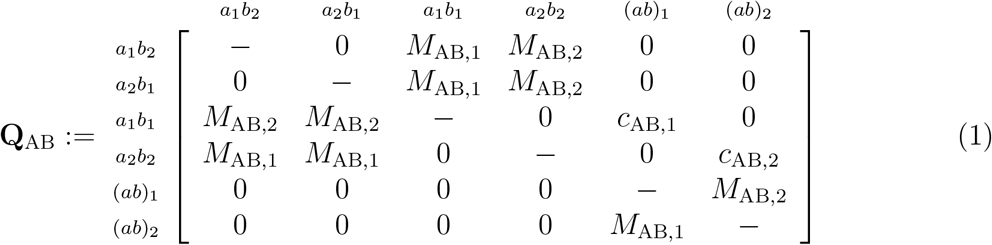

In our notation, *a*_*i*_*b*_*j*_ indicates that the lineage (denoted *a*) from species A is currently in subpopulation *i* and the lineage (denoted *b*) from species B is currently in subpopulation *j*, where *i, j* ∈ {1, 2}. Further, (*ab*)_*i*_ indicates that lineages *a* and *b* have coalesced and are in subpopulation *i*, where *i* ∈ {1, 2}. The coalescence rate in subpopulation *i* is *c*_AB,*i*_ = 2/*θ*_AB,*i*_, and the dashes along the diagonal represent the negative sum of the elements of the corresponding row, such that each row sums to zero (Kingman 1982). One can see that this matrix embeds several assumptions. First, we only allow for one event at any given instant. For example, we do not permit a lineage in one subpopulation and a lineage in another subpopulation to migrate simultaneously. Additionally, once lineages coalesce, they cannot uncoalesce. Under our model, at time *τ*_1_, lineages from species A are in subpopulation 1, and lineages from species B are in subpopulation 2.

The interval [*τ*_2_, ∞) is described using one of two possible instantaneous rate matrices. If the first coalescence is yet to occur, then the 20 × 20 matrix **Q**_ABC_ governs the dynamics. In the interest of space, this matrix is left to Appendix Figure S1. We should note, however, that because any two lineages are equally likely to coalesce in states when all three lineages are in subpopulation *i*, the rate of coalescence in subpopulation *i* in these scenarios is 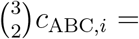 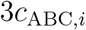, with *c*_ABC,*i*_ = 2/*θ*_ABC,*i*_. Under our model, at time *τ*_2_, the lineage *c* from species C must be in subpopulation 2, whereas lineages *a* and *b* from species A and B, or the coalesced lineage (*ab*), could be in either subpopulation. Above the root, once the first coalescence has occurred, the generic instantaneous rate matrix **Q**_*XY*_ models the dynamics. Letting *X* be a coalesced pair of lineages and *Y* be the remaining lineage, we have

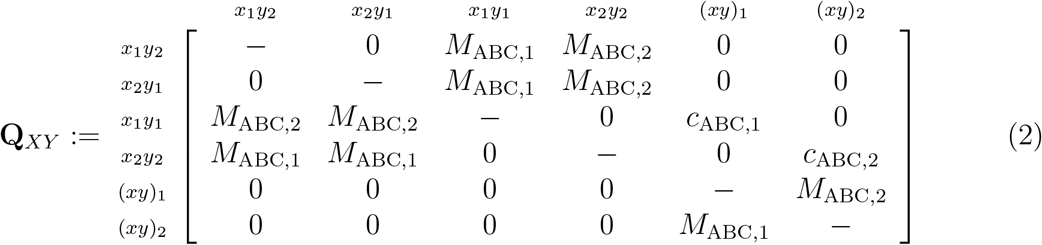

where *x* represents a lineage from coalesced pair *X* and *y* represents the lineage from species *Y*, and the same assumptions as for matrix **Q**_AB_ hold.

### Probability distributions of gene tree topologies under the model

We treat the dynamics in our model as a continuous-time Markov process that describes the waiting times between events. It follows that the waiting time to a given event is distributed exponentially with rate matrix **Q**, where events are independent as a consequence of the memoryless property of the exponential distribution (Ross 2014, Chapter 5).

Let *T*_1_ and *T*_2_ be random variables denoting the times to the first and second coalescence going backward in time, respectively. Further, let *G* be a random variable denoting the gene tree topology. With the set of parameters **Ω** = {*σ*, ***τ***, **M**, ***θ***}, let *f*_1_(*t*_1_, *t*_2_, *g* | **Ω**) be the joint density of gene tree *G* = *g* and coalescence times *T*_1_ = *t*_1_ and *T*_2_ = *t*_2_ when 0 ≤ *τ*_1_ ≤ *t*_1_ < *τ*_2_ ≤ *t*_2_. Further, let *f*_*j*_(*t*_1_, *t*_2_, *g*_*j*_ | **Ω**), *j* ∈ {2, 3, 4}, be the joint density of gene tree *G* = *g*_*j*_ and coalescence times *T*_1_ = *t*_1_ and *T*_2_ = *t*_2_ when 0 ≤ *τ*_1_ ≤ *τ*_2_ ≤ *t*_1_ < *t*_2_, where *g*_2_ = ((*ab*)*c*), *g*_3_ = ((*bc*)*a*), and *g*_4_ = ((*ac*)*b*). Note that *f*_1_(*t*_1_, *t*_2_, *g* | **Ω**) does not accumulate mass for *G* ≠ ((*ab*)*c*) because only lineages from species A and B can coalesce along the internal branch [*τ*_1_, *τ*_2_).

We can split the joint probability densities into a product of marginal densities *h*_AB,*ij*_ and *h*_ABC,*ij*_ of going from *i* lineages to *j* lineages in the internal branch and the branch above the root, respectively, because the waiting times to coalescence events are independent. Let *c*_AB,*s*_ and *c*_ABC,*s*_ be the coalescence rates leading to state *s*. For example, if the system has *n* lineages in subpopulation *i*, and a coalescence is about to occur there, then the rate of coalescence is 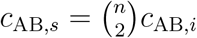 for the internal branch and 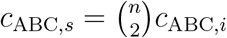 above the root. These densities are

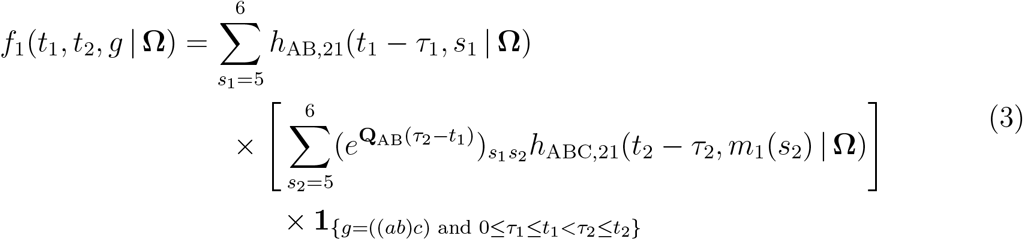

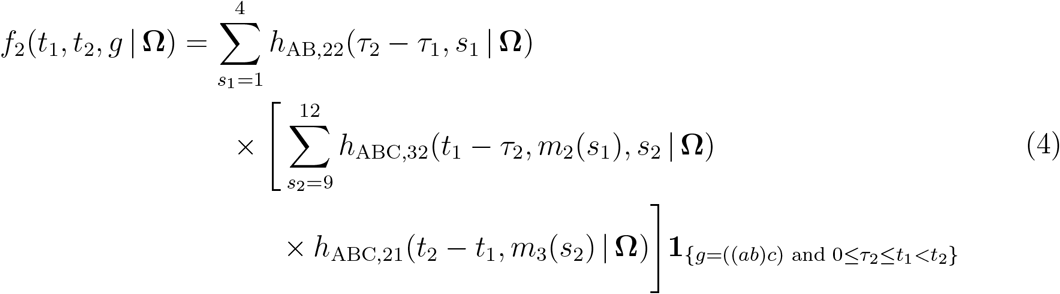

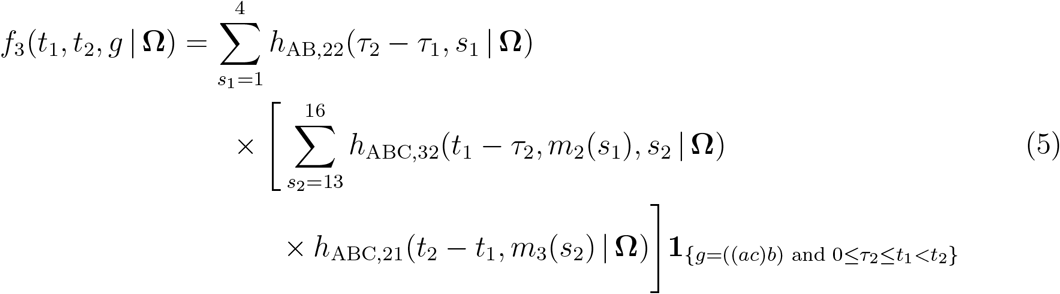

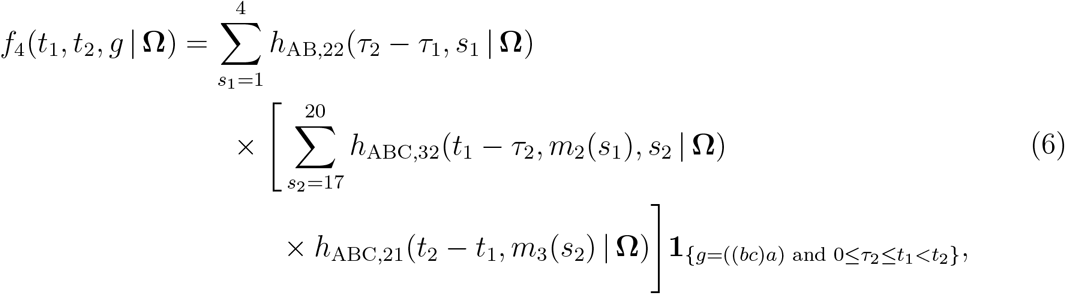

where

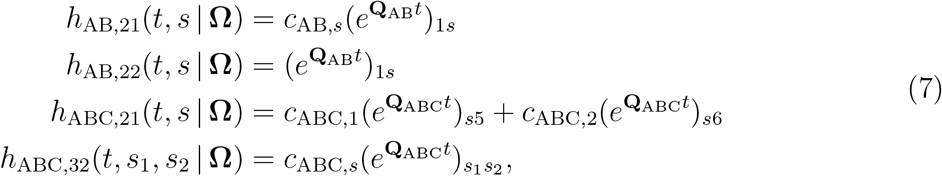

and

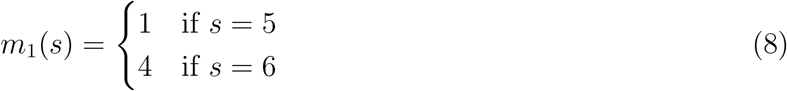

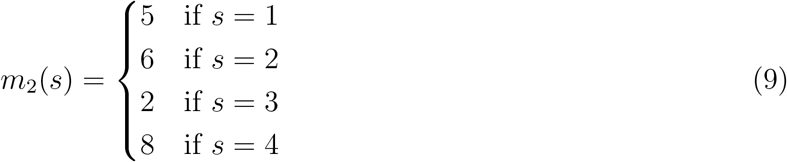

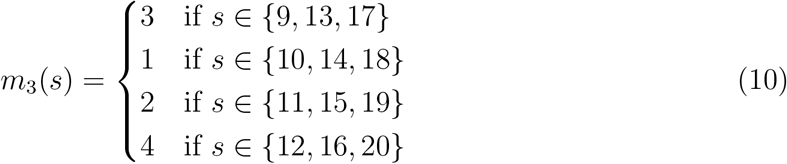

For *h*_AB,21_(*t, s* | **Ω**) and *h*_AB,22_(*t, s* | **Ω**), *s* represents the state of the system at time *t*. Specifically, *s* ∈ {5, 6} for *h*_AB,21_ and *s* ∈ {1, 2, 3, 4} for *h*_AB,22_. For *h*_ABC,32_(*t, s*_1_, *s*_2_ | **Ω**), *s*_1_ and *s*_2_ represent the states at time 0 and at time *t*, respectively. That is *s*_1_ ∈ {5, 6, 7, 8} and *s*_2_ ∈ {9, 10, …, 20}. Similarly, for *h*_ABC,21_(*t, s* | **Ω**), *s* represents the state of the system at *t* = 0, with *s* ∈ {1, 2, 3, 4}. The function *m*_1_(*s*) maps a coalesced state *s* in matrix **Q**_AB_ to a corresponding state in matrix **Q**_*XY*_ (*e.g.*, mapping state 5 in **Q**_AB_, where lineages *a* and *b* are coalesced in subpopulation 1, into corresponding state 1 of **Q**_*XY*_, where the coalesced lineage is in subpopulation 1 and lineage *c* is in subpopulation 2). The function *m*_2_(*s*) maps uncoalesced state *s* in matrix **Q**_AB_ to a corresponding state in matrix **Q**_ABC_ (*e.g.*, mapping state 4 in **Q**_AB_, where lineages *a* and *b* are both in subpopulation 2, to state 8 in **Q**_ABC_, where all uncoalesced lineages are in subpopulation 2). The function *m*_3_(*s*) maps a coalesced state *s* in matrix **Q**_ABC_ to a corresponding state in matrix **Q**_*XY*_ (*e.g.*, mapping state 19 in **Q**_ABC_, where lineages *b* and *c* are coalesced in subpopulation 2 and lineage *a* is in subpopulation 1, to state 2 in **Q**_*XY*_, where the coalesced lineage is in subpopulation 2 while the remaining lineage is in subpopulation 1). Though long, Equations 3–6 are simply capturing the probability density of all coalescent histories (Fig. 1b).

It follows that the probability of observing gene tree topology *G* given the model parameters **Ω** can be computed from these densities as

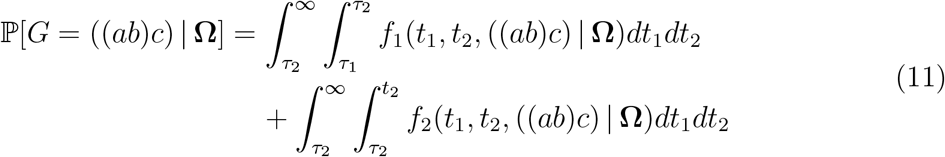

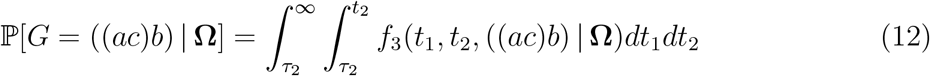

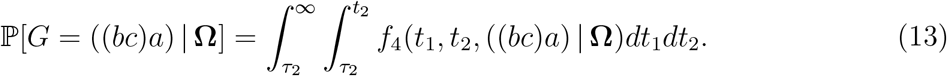

### Maximum likelihood parameter estimation

It is possible to compute the likelihood from a set of alignments {*A*_1_, *A*_2_, …, *A*_*K*_} at *K* independent loci as

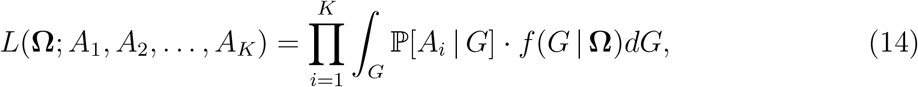

where ℙ[*A*_*i*_ | *G*] is the probability of observing the *i*th alignment *A*_*i*_ given gene tree *G*. Although the likelihood is tractable, it is computationally challenging and changes with the choice of substitution model used to compute the term ℙ[*A*_*i*_ | *G*] (Hey and Nielsen 2007). As such, it is common practice in likelihood methods to use estimated gene trees, 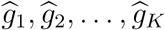, as input data, and treat these data as known. Assuming the collection of *K* inferred gene trees are independent, the likelihood of the model may be computed as

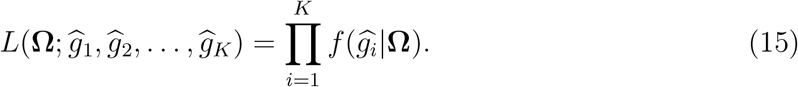

Denoting **Ω**_−*σ*_ = **Ω** \ {*σ*} as the parameter set **Ω** excluding the species tree topology *σ*, then for a fixed configuration of species tree topology *σ*, we search over all possible values of unknown parameters in **Ω**_−*σ*_ to maximize the log likelihood. Thus, each configuration of the species tree topology is linked with its own set of maximum likelihood estimates 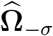. We infer the species tree and associated parameter estimates 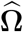 that give rise to the largest associated likelihood.

It must be emphasized that in this scenario of ancestral structure with two subpopulations, while the three possible species tree topologies are ((AB)C), ((AC)B), and ((BC)A), there are six total species tree model configurations in which species branch from ancestral subpopulations. The reason is that for each species tree topology ((*XY*)*Z*) relating species *X*, *Y*, and *Z*, it is possible that either *Y* and *Z* or *X* and *Z* descend from the same ancestral subpopulation. Therefore, we must find the maximum likelihood parameter estimates for six configurations of the species tree.

### Extension to *n* taxa

With three species, there are only three species tree topologies and two configurations of each topology to consider. However, as the number of taxa grows, the number of available topologies, which grows in complexity at a rate proportional to the double factorial of the number of taxa *n* (Felsenstein 2004), makes direct application of our method computationally infeasible. However, as illustrated in Figure 2, dividing the *n*-taxon problem into multiple sub-problems can ameliorate this burden. Because a rooted *n*-taxon tree is characterized by its set of rooted triples (Steel 1992), we can implement a supertree approach (*e.g*., DeGiorgio and Degnan 2010) to estimate the *n*-taxon species tree topology. This tactic reduces computation time to the order of 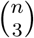 times the original complexity of the problem on three taxa.

**Figure 2:**
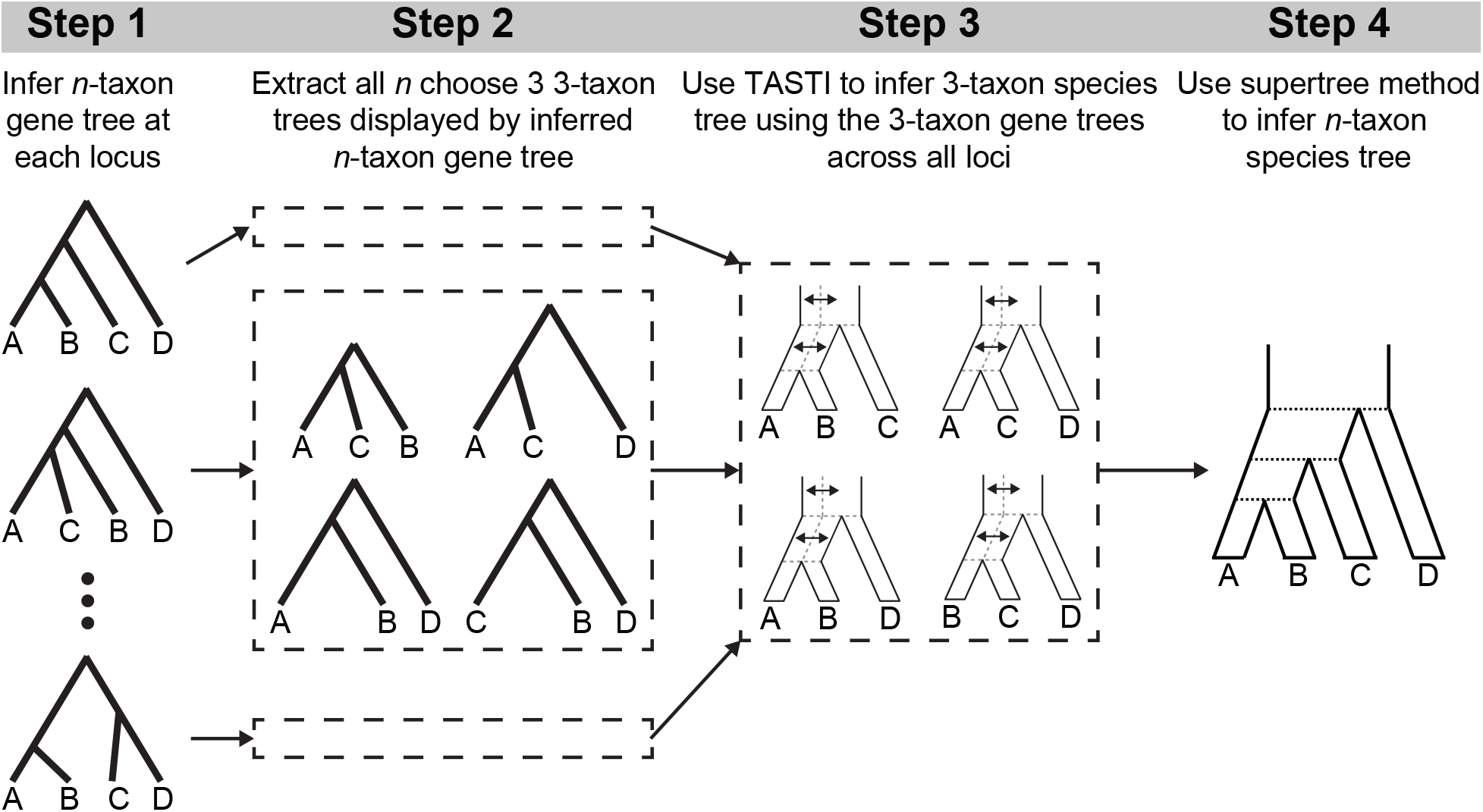
Schematic of a supertree approach for inferring *n*-taxon species tree topologies under a model of ancestral population structure. (Step 1) At each locus, infer a rooted *n*-taxon (here *n* = 4) gene tree. (Step 2) For each gene tree, extract the set of 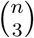 (here equaling four) rooted three-taxon gene trees compatible with each *n*-taxon gene tree. (Step 3) For a given set of three taxa (*e.g*., species A, B, and C), apply TASTI to infer a rooted three-taxon species tree under a model of ancestral population structure by using all three-taxon gene trees with the given set of taxa across all loci. (Step 4) Given the set of 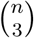 (here equaling four) rooted three-taxon species trees inferred by TASTI, use a supertree approach to infer a rooted *n*-taxon species tree topology for the full set of *n* taxa.

### Implementation

TASTI jointly optimizes over divergence times and migration rates, though the computation varies depending on the type of input data—*i.e.*, gene tree topologies with, or without, branch lengths. From this point forward, for simplicity and clarity, we drop the “hats” on our estimates for gene trees, as we treat them as fully known.

We first consider the case when branch lengths are included to make inference. Under this scenario, estimated coalescence times *t*_1_ and *t*_2_ are already fixed, eliminating the need to integrate out their values and permitting direct computation of the probability density given **Ω**. For coalescence times *t*_1*i*_ and *t*_2*i*_ of gene tree *g*_*i*_, computing the likelihood reduces to

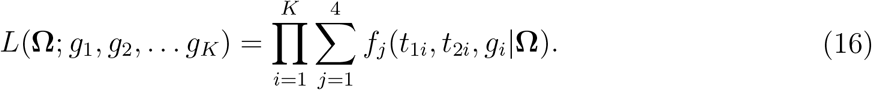

On the other hand, when simply inputting gene tree topologies, we can decrease the size of the parameter space by optimizing over the length of the internal branch, rather than estimating each divergence time separately, because our model assumes no coalescence occurs in the interval [0, *τ*_1_). The problem posed in Equations 11–13 also reduces to integrating only over *t*_1_, as the coalescence event at *t*_1_ uniquely determines the topology of the tree. Given data **y**= (*n*_*ab*_, *n*_*ac*_, *n*_*bc*_), where *n*_*xy*_ represents the number of three-taxon gene tree topologies in a sample displaying clade {*x, y*} for lineages *x* and *y* from species *X* and *Y*, we can compute the likelihood under **Ω** as

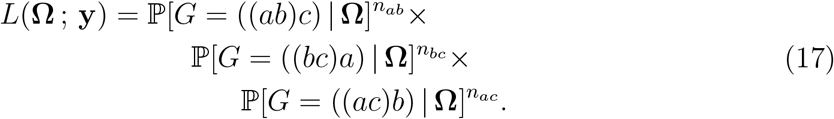

To implement a constrained optimization procedure, we first need to derive bounds on our unknowns. First, we consider the migration rate. Recall that the migration rate is given by *M* = 4*Nm*/*θ*, where *m* ∈ (0, 1]. This formulation gives rise to a lower bound of *M* > 0 when *m* is arbitrarily close to 0, since *m* = 0 would imply no migration between subpopulations and thus would prevent the process from stopping. Similarly, we have an upper bound on *M* of 4*N*/*θ* when *m* = 1. As the migration rate grows toward its upper bound, the species ancestry becomes effectively unstructured, generalizing our model to the standard multispecies coalescent.

If branch lengths are used as input, then the estimated coalescence times determine the bounds on the speciation times. Let {*T*_11_, *T*_12_, …, *T*_1*K*_} and {*T*_21_, *T*_22_, …, *T*_2*K*_} be the collections of first and second coalescent times at *K* loci, respectively, going backward in time. Then by our model construction, *τ*_1_ ∈ [0, min{*T*_11_, *T*_12_, …, *T*_1*K*_}] and *τ*_2_ ∈ [*τ*_1_, min *T*_21_, *T*_22_, … .*T*_2*K*_]. In a maximum likelihood framework, if branches of a gene tree in the sample are uninformative, it is then possible to encounter a “star tree” scenario in which *τ*_1_ = *τ*_2_ = 0. Here, we cannot resolve the relationship among the taxa and are left to infer the unresolved species tree topology (ABC). Additional details regarding the implementation of the optimization procedure are discussed in the Appendix under section *Parameter estimation for gene trees with branch lengths*.

When only topologies are considered, however, we simply need to bound the length of the internal branch of the species tree. To do this, we claim it is reasonable to search only over internal branch lengths that allow for some reasonable level of gene tree discordance (Pamilo and Nei 1988; Hudson 1983; Tajima 1983). This is because as the length of the internal branch of the species tree increases, the probability of discordance in the presence of population structure, even with low migration rates, goes to zero. Letting *τ* represent the length of the internal branch, this probability of discordance is computed by

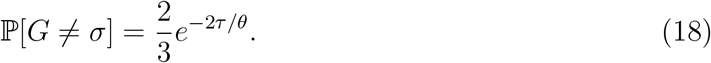

We propose that the lowest reasonable level of discordance to assume is ℙ[*G* ≠ *σ*] = 0.05, implying that out of 100 sampled loci, only five will exhibit discordance. Imposing this bound is similar in spirit to other methods which assume a minimum level of incomplete lineage sorting (Than et al. 2008; Wu 2012).

## Results

In this section we aim to examine the performance and robustness of our proposed method using various forms of input data, and compare this performance to existing likelihood-based methods MP-EST (Liu et al. 2010b), STELLS2 (Pei and Wu 2017), and STEM2.0 (Kubatko et al. 2009). MP-EST is a pseudo-likelihood approach based on triples of taxa, whereas STELLS2 and STEM are likelihood approaches based on gene tree topologies and gene tree topologies with branch lengths, respectively. For TASTI, we considered using gene tree topologies with and without branch lengths in cases where the input data are both inferred and known with certainty. The gene trees in our simulations were generated using the coalescent simulator *ms* (Hudson 2002). To simulate sequence alignments conditional on these gene trees, we employed Seq-Gen (Rambaut and Grass 1997) to generate one kilobase (kb) and 0.5 kb long sequences (with an additional outgroup sequence) under the HKY substitution model (Hasegawa et al. 1985) with a transition-transversion ratio of 4.6 and base frequencies A, T, C, and G respectively equal to 0.3, 0.3, 0.2, and 0.2.

From these simulated sequences, we used dnamlk of PHYLIP under the HKY substitution model (Felsenstein 1989) to infer gene trees using maximum likelihood assuming a molecular clock with a transition-transversion ratio of 4.6 and empirical base frequencies. We applied this pipeline to create samples of 100 replicates of *K* independent loci, where *K* ranged from 10 to 10^4^, under fixed species tree topology ((AB)C). We let *M*_AB,1_ = *M*_AB,2_ = *M*_ABC,1_ = *M*_ABC,2_ = *M* and *θ*_AB,1_ = *θ*_AB,2_ = *θ*_ABC,1_ = *θ*_ABC,2_ = *θ*. We set a constant effective population size of *N* = 5 × 10^4^ across both subpopulations, and employed a per-site per-generation mutation rate of *μ* = 2.5 × 10^−8^, yielding *θ* = 0.005. When gene tree topologies were used as input, this resulted in a bound on the length of the internal branch of the species tree of *τ*_2_ − *τ*_1_ ≈ 6.5 × 10^−3^, as derived from Equation 18. These parameter settings were inspired by the great ape dataset of Burgess and Yang (2008).

### Ancestral population structure alters expected gene tree distributions

While symmetries among the distribution of gene tree topologies are expected under the standard multispecies coalescent (Allman et al. 2011), population structure skews this distribution, as lineages above the root are no longer guaranteed to have equal probability of coalescing (Slatkin and Pollack 2008). In Figure 3, distributions of gene trees are plotted for varying levels of migration. Indeed, as *M* → 4*N*/*θ*—the scenario under which our model reduces to the standard multispecies coalescent—this symmetry between topologies ((*bc*)*a*) and ((*ac*)*b*) is present. However, as the migration rate decreases, the distribution skews toward a dominant ((*bc*)*a*) topology, and symmetry between gene tree topologies ((*bc*)*a*) and ((*ac*)*b*) no longer persists. Further, while estimated gene tree topologies (Fig. 3c) obey the same large-sample distribution as the topologies known exactly (Fig. 3a), there is an increased variability in their distribution across samples, which in general is detrimental to inference (Casella and Berger 2002, Chapter 10).

**Figure 3:**
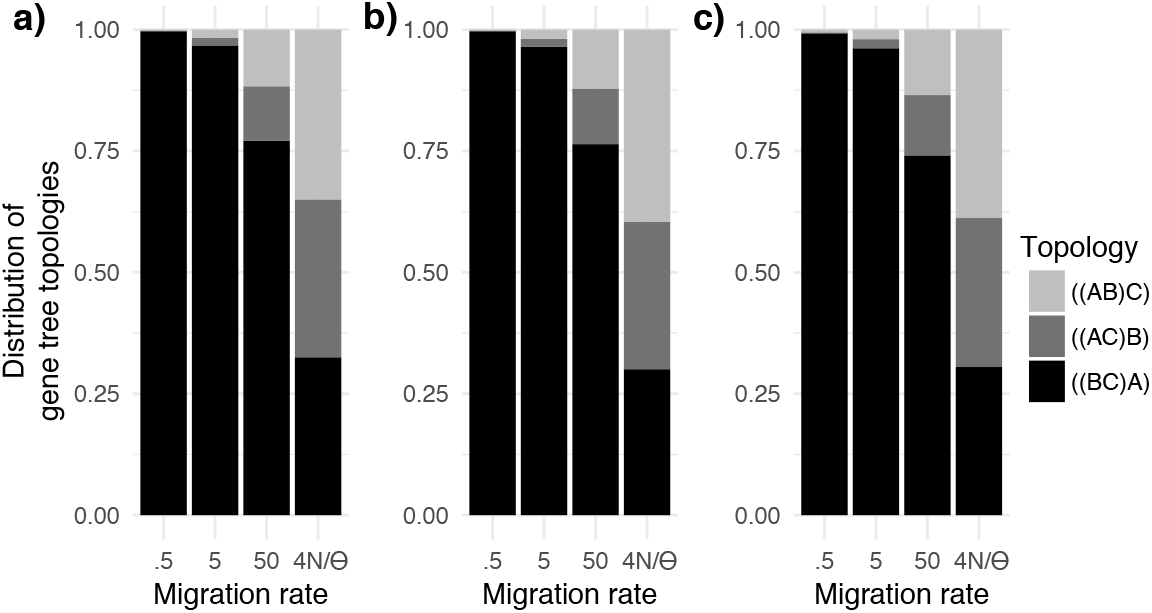
Distributions of gene tree topologies over various rates of migration for data. *a*) Exact distributions of gene trees under the model using parameter settings given at the beginning of the *Results* section. *b*) Distributions of gene trees under the model from 10^3^ simulated gene trees. *c*) Distributions of gene trees under the model from 10^3^ gene trees estimated from sequence of length one kb using the approach detailed at the start of the *Results* section.

TASTI incorporates the possibility for these skewed distributions of gene tree topologies. In what follows, we demonstrate that TASTI provides reasonable estimates of the true species tree topology under conditions in which other methods have previously failed (DeGiorgio and Rosenberg 2016). We simulated data as described at the start of this section, assuming a species tree with speciation times *τ*_1_ = 2.5 × 10^−3^ and *τ*_2_ = 2.75 × 10^−3^ mutation units. This setting, with such a short internal branch, makes correct inference especially challenging. In order to understand the effects of various evolutionary parameters on TASTI’s performance, we first conduct a detailed exploration of the method’s accuracy. A summary of TASTI’s accuracy is in Figure 4, while tables of median parameter estimates and 95% confidence intervals are in Tables S1 and S2, respectively. Summaries of additional simulations studying the performance of TASTI against MP-EST, STELLS2, and STEM2.0 under a variety of different simulation settings are in Figure 5.

**Figure 4:**
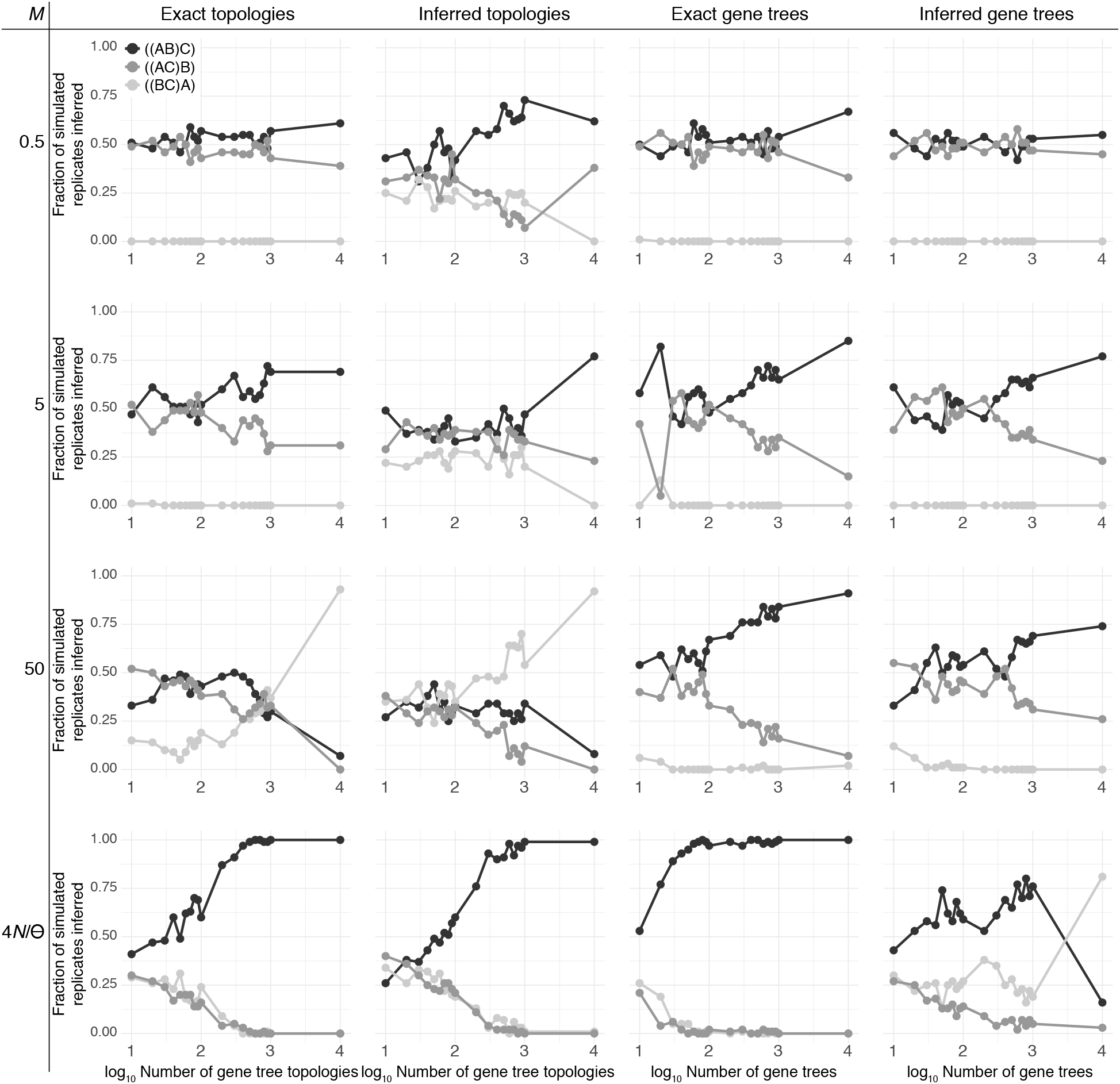
Accuracy of the maximum likelihood estimator of *σ* as a function of the number of input gene trees. Accuracy is based on the proportion of 100 simulated replicates of *K* loci (before filtering “star trees”), with *K* ranging from 10 to 10^4^, where our method inferred a specific species tree topology under a scenario with *τ*_1_ = 2.5 × 10^−3^ and *τ*_2_ = 2.75 × 10^−3^.

**Figure 5:**
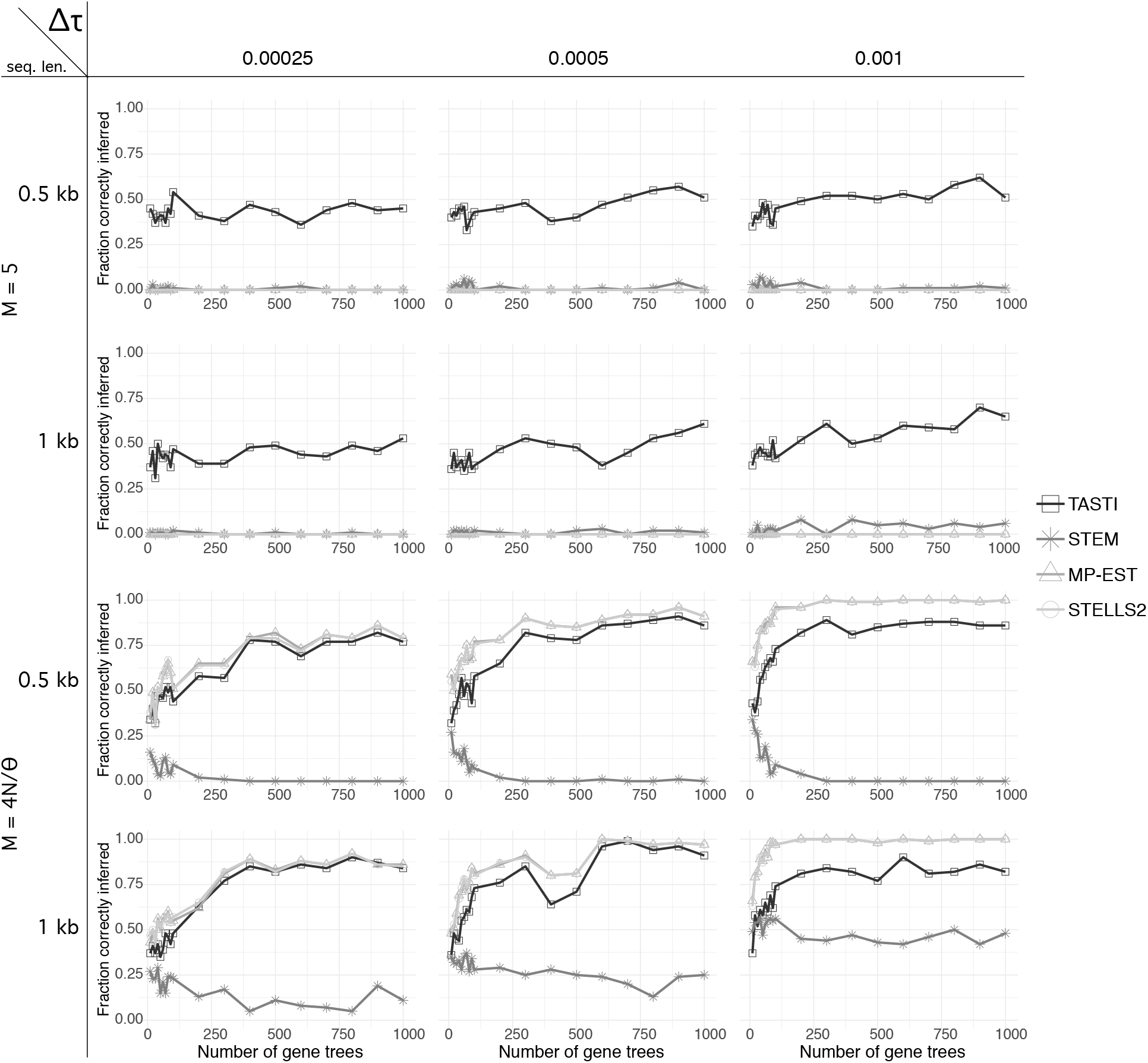
Accuracy of TASTI, MP-EST, STELLS2, and STEM2.0 when estimating the true species tree topology *σ* as a function of the number of input gene trees. Accuracy is based on the proportion of 100 simulated replicates of *K* loci with *K* ranging from 10 to 10^3^. In these simulations, we only use inferred gene tree topologies as input to TASTI. Comparisons are made across three internal branch lengths (Δ*τ* = *τ*_2_ − *τ*_1_) in cases when populations are both structured and unstructured and when gene trees are estimated from both one and 0.5 kb sequences.

Our results drive home several key points. First, phylogenetic inference methods are sensitive to the quality of input data, and TASTI taking gene tree topologies as input tends to be more robust to stochasticity encountered in practice than when gene tree branch length information is incorporated into TASTI’s input as well. However, if accurate data on gene tree branch lengths are available, then TASTI will naturally perform better when this information is included rather than obscured. This is especially true when multiple candidate species trees are equally likely under a model of gene tree topologies alone. Furthermore, when species trees have longer internal branch lengths, then more accurate input data can be estimated, yielding more reliable downstream inference of the complete species trees. Lastly, while TASTI is competitive with alternative methods under the standard multispecies coalescent, it is the only method that exhibits favorable performance in the presence of ancestral population structure.

### Gene tree topology data permits accurate, robust species tree inference

The first column of Figure 4 shows the performance of TASTI with exact gene tree topologies for data input across all investigated levels of migration. In the case of no ancestral structure (*M* = 4*N*/*θ*), we obtain consistent estimates of the true species tree even with a small number of sampled loci. Similarly, when the ancestral populations are structured with *M* = 5, TASTI’s species tree estimate begins favoring the truth over alternative topologies fairly quickly. In contrast, with a combination of a higher rate of migration between subpopulations (*M* = 50) and a short internal branch of the species tree, the effects of ancestral population structure begin to break down. Consequently, the distribution of gene tree topologies resembles one we would expect under the standard multispecies coalescent with sister species B and C, such that TASTI is lead to infer the dominant gene tree topology ((BC)A) as the true species tree. When migration is at its lowest (*M* = 0.5), and especially with few sampled loci, the data do not convey any detectable signal in the distribution of topologies. That is, nearly all of the observed topologies are of the form ((*bc*)*a*), and the true species tree is difficult to determine. Interestingly, though ((*bc*)*a*) is by far the dominant gene tree in these input data, ((BC)A) is never TASTI’s estimated maximum likelihood species tree. This result is expounded upon in Appendix section *Bounding the length of the internal branch of the species tree in the low signal setting*.

The second column of Figure 4 illustrates TASTI’s accuracy when gene tree topologies are no longer known with certainty, but rather, are inferred from one kb sequences. A comparison of these results with those in the first column suggests that TASTI is robust to noise incurred in the process of gene tree inference. While accuracy in general is reduced, the overall trends are still present. For the no structure setting, and when ancestral populations are structured with *M* = 5, TASTI still favors the true species tree topology ((AB)C), but with less immediacy and certainty as is found in the case when topologies are known exactly. When *M* = 50, TASTI goes from slowly favoring the misleading species tree topology ((BC)A) to quickly being positively misleading for ((BC)A). Interestingly, when migration is at its lowest, the added noise from inferred topologies actually improves TASTI’s accuracy. This is most likely because with additional noise we observe topology ((*ab*)*c*) more frequently than when gene trees are known exactly. Thus, we more regularly infer *σ* = ((AB)C).

### Knowledge of gene tree branch lengths improves estimation

Incorporating branch length data from input gene trees can improve inference under TASTI, as knowledge about presence or absence of symmetries in the distribution of gene tree topologies may sometimes be insufficient for making the correct judgment. The performance of TASTI when gene tree topologies with branch lengths are known with certainty, displayed in the third column of Figure 4, provides a benchmark best case that TASTI can hope to achieve under our highly challenging simulation settings. In the case when ancestral populations are not structured we see perfect performance by TASTI with few input loci. With ancestral structure, we still achieve good performance under each level of migration, with weaker trends as the level of structure between subpopulations increases. Remarkably, while TASTI’s estimate of the species tree topology is inconsistent when *M* = 50, the addition of branch lengths rescues inference. Another advantage of using branch lengths is that we eliminate the need to set an upper bound on the length of the internal branch heuristically, as the estimated gene trees determine this automatically.

Using estimated gene trees and branch lengths introduces increased variability into the model when compared to the noise introduced by only using estimated topologies. In general, while we obtain fairly accurate species tree estimates using gene tree data known with certainty, the overall trend is that TASTI experiences a larger reduction in accuracy from using estimated gene trees than it does from using estimated topologies alone. For example, we can see that under the standard multispecies coalescent, though we obtained near-perfect performance when branch lengths were known with certainty, the variability in the estimated branch lengths greatly reduces TASTI’s accuracy. This trend is apparent across all migration rates, but it increases in severity as the level of structure decreases. This is consequent to the claim that if gene tree branch lengths are shorter, they are more difficult to estimate accurately (DeGiorgio and Degnan 2014). This notion is explored more deeply in the following section.

### Species trees with longer total branch length permit better inference

Our simulations test the accuracy of TASTI under a highly challenging scenario where the internal branch of the species tree is excessively short. With a short internal branch under ancestral population structure, the chance of observing gene trees concordant with the species tree decreases. Additionally, for a fixed migration rate and divergence time *τ*_1_, the time to the MRCA among lineages in the phylogeny decreases on average with a decreasing internal branch length. These shorter gene tree branch lengths make accurate gene tree inference more difficult, as loci become less informative because fewer mutations will have occurred among the species (DeGiorgio and Degnan 2014).

The impact of the short gene tree branch lengths can be seen directly by comparing the accuracy of TASTI when applied to gene trees known with certainty as opposed to when it is applied to inferred gene trees (Fig. 4, third and fourth columns). When migration is low, gene tree branch lengths are longer, and hence more informative. Thus, in lower-migration settings (*e.g*., when *M* = 0.5 or *M* = 5), we can infer more accurate gene trees and branch lengths than in higher-migration scenarios. However, when migration is high (*e.g*., when *M* = 50) or when the ancestral species are unstructured, gene tree branch lengths are much shorter, thereby leading to less accurately inferred divergence times. We can see that the accuracy of TASTI suffers the most when gene tree branch lengths are inferred under the no ancestral structure and high migration scenarios. Meanwhile, species tree inference is not dramatically affected when migration is low.

With this in mind, we tested the influence of noisy gene trees on TASTI’s performance in two ways. First, we considered the effect of less accurate input gene trees by following an identical simulation protocol to the one described above, with the exception that we inferred gene trees from 0.5 kb instead of one kb long regions. Results display similar, though slightly worsened, performance when compared to our original simulations (compare Figs. 4 and S2). We also tested the accuracy of TASTI with a longer internal branch of the true species tree, such that the time to the MRCA is on average longer. To evaluate this scenario, we simulated 100 replicates of 10^3^ loci using the same parameters and protocols as previously described, with the exception that we increased the age *τ*_2_ of the species tree root. Specifically, we employed divergence times of *τ*_1_ = 2.5 × 10^−3^ and *τ*_2_ = 5.25 × 10^−3^. As expected, we observe the same trends as in our original simulations (Fig. 4), but with overall more accurate species tree inference (Table 1). This is still a challenging setting, noting that the probabilities of observing concordant gene trees are 3.62 × 10^−3^, 3.46 × 10^−2^, 0.236, and 0.596 for *M* = 0.5, 5, 50, and 4*N*/*θ*, respectively, with the longer internal branch length. This can be compared against concordance probabilities of 1.83 × 10^−3^, 1.75 × 10^−2^, 0.122, and 0.366 for *M* = 0.5, 5, 50, and 4*N*/*θ*, respectively, with the shorter internal branch length.

**Table 1:** Accuracy of the maximum likelihood estimator as a function of migration rate *M* = 4*Nm*/*θ*. Accuracy is based on the percentage of 100 simulated replicates of 10^3^ loci that inferred a specific species tree topology under a scenario with *τ*_1_ = 2.5 × 10^−3^ and *τ*_2_ = 5.25 × 10^−3^.

| $M$ | Exact topologies | | | Inferred topologies | | | Exact gene trees | | | Inferred gene trees | | |
| --- | --- | --- | --- | --- | --- | --- | --- | --- | --- | --- | --- | --- |
|  | ((AB)C) | ((BC)A) | ((AC)B) | ((AB)C) | ((BC)A) | ((AC)B) | ((AB)C) | ((BC)A) | ((AC)B) | ((AB)C) | ((BC)A) | ((AC)B) |
| 0.5 | 60 | 23 | 17 | 63 | 19 | 18 | 84 | 0 | 16 | 75 | 0 | 25 |
| 5 | 73 | 27 | 0 | 75 | 25 | 0 | 94 | 0 | 6 | 93 | 0 | 7 |
| 50 | 77 | 23 | 0 | 75 | 25 | 0 | 94 | 0 | 6 | 91 | 0 | 9 |
| $4N/\theta$ | 74 | 13 | 13 | 52 | 25 | 23 | 100 | 0 | 0 | 84 | 5 | 11 |

While our study operated on gene trees inferred by maximum likelihood, the use of Bayesian methods to infer gene trees may alleviate these concerns by improving gene tree estimates and decreasing the chance of estimating gene trees with branches of length zero (DeGiorgio and Degnan 2014). Alternatively, it is possible that imposing a constraint on a minimum value on *τ*_2_ − *τ*_1_ when using whole gene trees would avoid the deleterious effects of gene trees with poorly estimated branch lengths.

### Comparison of competing methods in the three-taxon setting

We used simulations to compare TASTI to three other methods for inferring species trees from gene trees: MP-EST, STELLS2, and STEM2.0. We designed the study similarly to the one presented in Figure 4, examining the methods’ collective performance over varying internal branch lengths and levels of population structure. Specifically, we let (*τ*_2_ − *τ*_1_) ∈ {2.5 × 10^−4^, 0.5 × 10^−4^, 1.0 × 10^−3^} mutation units and *M* ∈ {5, 4*N*/*θ*}. We only investigate the more realistic case of estimated gene trees as input, where gene trees were inferred from 0.5 and one kb long sequences. For TASTI, we study the setting where gene tree topologies are used as input. Other details of the simulations, including parameters *N* and *θ*, remain the same as in previous simulations.

Results (Fig. 5) show the proportion of times out of 100 replicate simulations that each method inferred the correct species tree for the given parameter settings. Comparing between the top two rows, we see that TASTI’s accuracy increases with more accurate input topologies and with a longer internal branch, where the signals are stronger. Meanwhile, all other methods infer the wrong species tree topology nearly (or exactly) 100 percent of the time, demonstrating that MP-EST, STELLS2, and STEM2.0 are not robust to ancestral population structure.

The bottom two rows, on the other hand, evaluate all methods when the populations are unstructured. Here, MP-EST and STELLS2 show the same consistent performance; TASTI is competitive with these two methods. Though STEM is known to perform well when gene trees are known exactly, we see that this method is not robust to empirical data and therefore underperforms, especially when the gene trees are inferred from shorter sequences.

### Accuracy of the supertree approach applied to four taxa

With the wide availability of multilocus sequence data from a number of species (*e.g*., Johnson et al. 2013; Lin et al. 2014; Yang et al. 2015; Shen et al. 2016), it is common practice to construct species-level phylogenies for more than three taxa. However, when performing model-based species tree estimation, the inference problem becomes increasingly complex and computationally difficult (Liu and Pearl 2007; Drummond and Rambaut 2007; Heled and Drummond 2009; Wu 2012). We therefore chose to evaluate the performance of TASTI for an increased problem size. Using again the identical simulation protocol, we generated data assuming a four-taxon species tree with fixed species tree topology (((AB)C)D) and speciation times *τ*_1_ = 1.25 × 10^−2^, *τ*_2_ = 1.375 × 10^−2^, and *τ*_3_ = 1.5 × 10^−2^ mutation units. Species A originated from subpopulation 1 whereas species B, C, and D originated from subpopulation 2, as depicted in Figure 6. We again set a constant effective population size of *N* = 5 × 10^4^ across both of these subpopulations, and considered both the unstructured ancestral population case as well as the case with symmetric migration rate of *M* = 5 between subpopulations in ancestral species.

**Figure 6:**
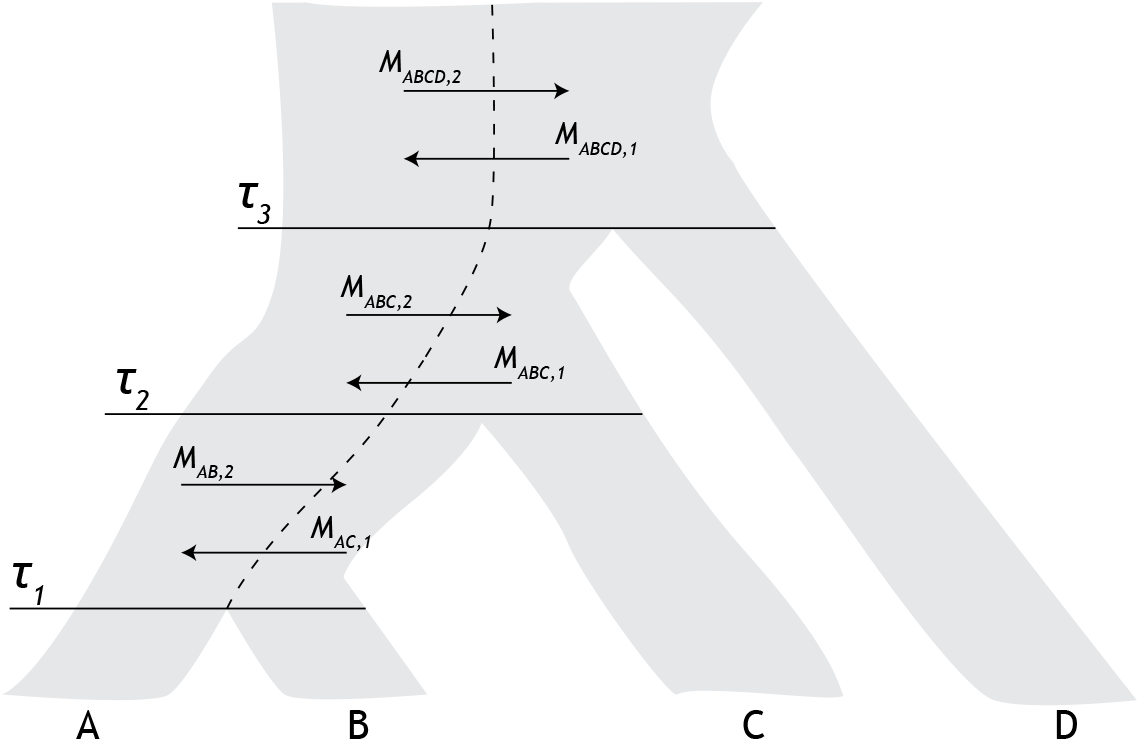
Model four-taxon species tree used in our supertree simulations, displaying the relationships among species A, B, C, and D, with divergence times *τ*_1_, *τ*_2_, and *τ*_3_. Ancestral species belong to one of two subpopulations with migration between subpopulations at rates *M*_AB,1_ and *M*_AB,2_ directly ancestral to species A and B, *M*_ABC,1_ and *M*_ABC,2_ directly ancestral to the split of species A and B with species C, and *M*_ABCD,1_ and *M*_ABCD,2_ above the root. Lineages from species A descend from subpopulation 1, whereas lineages from species B, C, and D descend from subpopulation 2. Under our simulation scenarios, we set *M*_*X,i*_ = *M*_*Y,j*_ = *M* for all species *X* and *Y* and for all subpopulations *i* and *j*.

To infer four-taxon species trees, we followed the procedure depicted in Figure 2. For each simulated replicate, we extracted the 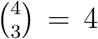 rooted triples displayed by each four-taxon gene tree at each of the *K* simulated loci. That is, at every locus, we extracted the rooted gene tree (topology with corresponding branch lengths) associated with each set of three species, such that we obtained rooted gene trees for the sets of taxa {*a, b, c*}, {*a, b, d*}, {*a, c, d*}, and {*b, c, d*}. For each set of three species, we estimated a three-taxon species tree topology using TASTI based upon the set of rooted triples for that set of three species across the *K* simulated loci. This procedure yielded 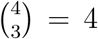 three-taxon species tree topology estimates. These four triples were then used to construct a supertree using the modified min cut supertree algorithm (Page 2002).

We chose to use the modified min cut supertree algorithm based on six properties it exhibits (Page 2002; Semple and Steel 2003). First, any species that is in one of the input gene trees is also in the inferred supertree. Second, if a tree exists that could display each input tree as a subtree, then the supertree will display each of these trees. Third, the order in which gene tree topologies are provided to the algorithm does not affect its supertree estimate. Fourth, changing the labeling of species in the set of input trees will produce the same output supertree with the set of relabeled species. Fifth, the algorithm runs in polynomial time as a function of the number of input species *n*. Finally, nestings that are not contradicted are also displayed in the supertree. Though we employ a specific supertree algorithm here, other supertree approaches may have complementary desirable properties that could lead to improvements in performance. However, an extensive assessment of the wide diversity of supertree algorithms (Wilkinson et al. 2005) is beyond the scope of this article.

In general, the most frequently inferred species tree topologies are the true topology (((AB)C)D) and the partially unresolved topology ((AB)CD) (Fig. S7). These unresolved trees occur as a consequence of the supertree construction algorithm when inferred triples are in conflict. We further investigated the performance of TASTI by examining the mean Robinson-Foulds (RF) distance (Robinson and Foulds 1981) between the inferred species tree topology and the true topology for each set of 100 replicates given *K* simulated loci. We followed the procedure of DeGiorgio and Degnan (2014) and defined the RF distance as the sum of the number of false positive clades and false negative clades between the inferred topology and the true topology, such that the distance metric applies to multifurcating trees. Mean RF distances, numbers of false positive clades, and numbers of false negative clades across all simulations are summarized in Figure S8. Results show that false negative clades occur more frequently than false positive clades. This result indicates that the largest errors of this approach are due to partially-unresolved species trees, rather than species trees displaying incorrect clades. Further, RF distance decreases as the number of input loci increases, with the exception of species tree estimates based on inferred gene trees with branch lengths.

### Comparing accuracy of multiple methods applied to four taxa

In order to place the above results in better context, we compared TASTI to MP-EST, STELLS2, and STEM2.0 in the four-taxon setting as well. We followed the same protocol for simulation that was used in the section *Accuracy with the supertree approach applied to four taxa*. In each simulation, we assumed equispaced divergence times (*e.g.*, *τ*_3_ − *τ*_2_ = *τ*_2_ − *τ*_1_) along an asymmetric species tree. The simulations include settings where the divergence times are separated by one of 2.5 × 10^−4^, 5.0 × 10^−4^, or 1.0 × 10^−3^ mutation units. We considered the structured case with migration rate *M* = 5 and the unstructured case. We applied this comparison only to inferred gene trees, where the gene trees were estimated from one and 0.5 kb sequences. For TASTI, we only used gene tree topologies as input.

The results from this simulation, summarized using the RF distance in Figure 7, behave as expected (the same simulations, described by number of false positive clades and false negative clades, are reported in Figures S5 and S6, respectively). The top two rows, showing simulations when the populations are structured, indicate that MP-EST, STELLS2, and STEM2.0 behave similarly poorly; each method on average infers a tree that is as far away in RF distance from the true tree as possible. TASTI, on the other hand gets closer to the true tree, in particular as the internal branches grow longer. In the unstructured case, while TASTI, MP-EST and STELLS2 are all competitive, STELLS2 uniformly performs the best. TASTI and MP-EST perform very similary because MP-EST takes a pseudo-likelihood approach that, like TASTI, is also based in rooted triples. Because STEM2.0 is sensitive to noisy data, this method underperforms relative to the other three.

**Figure 7:**
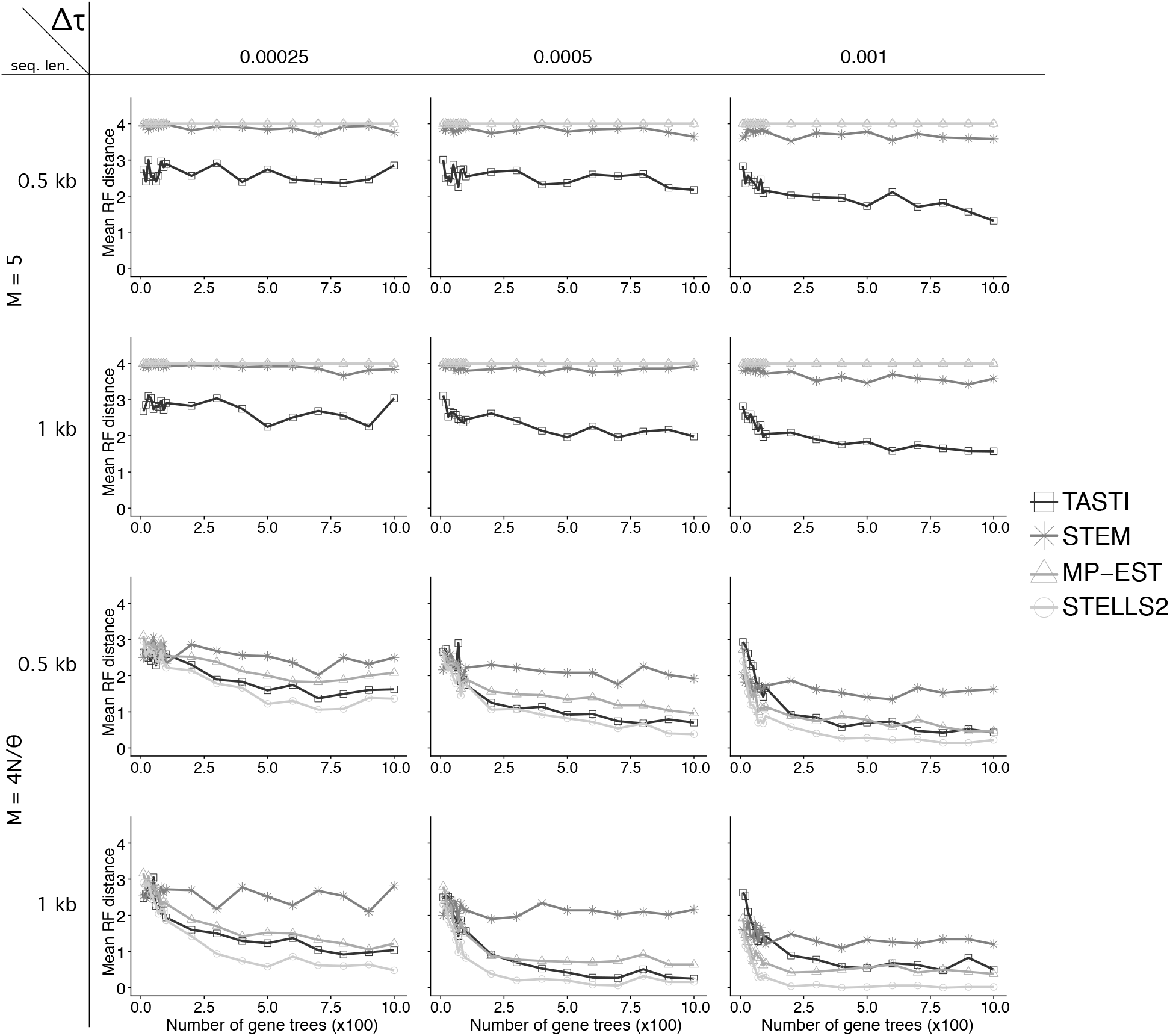
Accuracy of the competing methods’ estimators of *σ* as a function of the number of input gene trees. Accuracy is based on the average Robinson-Foulds (RF) distance between the estimated and true species trees across 100 simulated replicated of *K* loci, with *K* ranging from 10 to 10^3^. We considered a migration rate *M* = 5 as well as the unstructured scenario.

### Application to data from the Anopheles gambiae complex

The *Anopheles gambiae* complex is comprised of several morphologically indistinguishable yet genetically different species of mosquitos living in sympatry in sub-Saharan Africa (Davidson 1962). Though the species belonging to the *Anopheles gambiae* complex are closely related, only a small fraction of them possess significant malaria vectorial capacity. Studying their ancestry is critical as it may lead to key insights regarding how evolutionary history affects the development of traits that drive successful malaria vectors (Neafsey et al. 2015). Fontaine et al. (2015) originally studied six species in this complex: *Anopheles arabiensis*, *Anopheles coluzzii*, *Anopheles gambiae sensu stricto*, *Anopheles melus*, *Anopheles merus*, and *Anopheles quadriannulatus*. In their work, they identified that two of the most important malaria vectors in the complex, *An. arabiensis* and *An. gambiae*, are in fact not the most closely related. However, the similarities in their genome could be attributed to introgression between them found on their autosomes. Wen et al. (2016a) built on the analyses of Fontaine et al. (2015) by applying a phylogenetic network model to the *Anopheles* data. Given that the phylogenetic signals found in this dataset are more complex than one expects to find assuming incomplete lineage sorting alone, we deemed this a viable system in which to apply TASTI. In what follows, we conduct analyses of the autosomes for the *Anopheles gambiae* complex, using a seventh species, *An. christyi*, as an outgroup.

A single individual from each of the six species was sequenced at high coverage. Wen et al. (2016a) subset these high-coverage samples to generate independent genomic regions. Because TASTI also assumes independent loci, we used these alignments as input. After subsetting by Wen et al. (2016a), an ample 2,791 independent alignments across the autosomes with an estimated mean length of 3.4 kb remained in this dataset (Figure S1 of Wen et al. (2016a) shows a histogram of locus lengths). Following the procedures of both Fontaine et al. (2015) and Wen et al. (2016a), we estimated maximum likelihood gene tree topologies under the GTR+Gamma model at each locus using RAxML (Stamatakis 2014). Figure S9 displays the distributions of input rooted triples estimated in this manner, revealing asymmetries among some of the distributions of topologies. Following Wen et al. (2016a), we also generated 100 bootstrap alignments at each locus and estimated the maximum likelihood topology for each bootstrap replicate. These bootstraps can be used to account for uncertainty in gene tree topology estimates, analogous to the Bayesian approach described by Yu et al. (2012). Briefly, let *K* be the number of loci, and for each locus *k*, let *g*_*k1*_, *g*_*k2*_, …, *g*_*kq*_ be the *q* different gene tree topologies estimated at that locus from the bootstrap alignments. Denote *α*_*k*1_, *α*_*k*2_, …, *α*_*kq*_ as the proportion of times that gene tree topologies 1, 2, …, *q* were respectively inferred from these alignments at locus *k*, such that 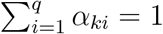. Then, letting 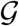 be the set of all gene tree topologies computed across the bootstrap replicates across *K* loci, we define for each 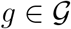 the sum of bootstrap probabilities associated with all loci whose topology is g as pg. That is, for each 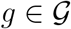, we have 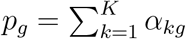 such that 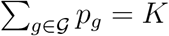. Let 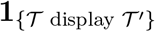 be an indicator random variable that takes the value 1 if tree topology 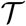 displays tree topology 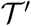, and 0 otherwise. We thus modify the likelihood of a species tree in Equation 17 to be

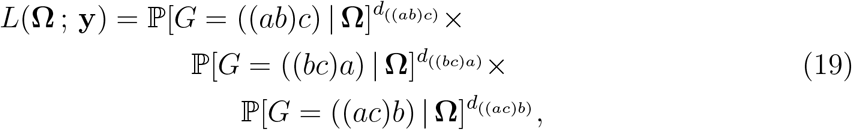

where

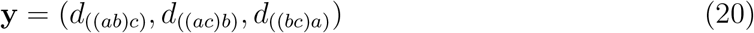

and

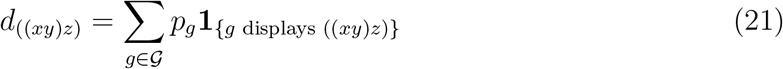

denotes the sum of probabilities of gene trees displaying the rooted triple ((*xy*)*z*), such that *d*_((*xy*)*z*)_ + *d*_((*xz*)*y*)_ + *d*_((*yz*)*x*)_ = *K*.

To conduct our analyses, we first needed to obtain an estimate for the diversity parameter *θ* = 4*Nμ*. Fontaine et al. (2015) sequenced several individuals from each of the six species under study in the complex at lower coverage. We computed the mean pair-wise sequence difference 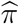 (Tajima 1983) within populations after filtering out indels using VCFtools (Danecek et al. 2011). Recalling that 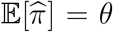 (Tajima 1983), we proceeded by using 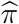 as an estimate of *θ*. Across the genome, however, the estimate of *θ* within each species in the complex varied by over an order of magnitude (from ~ 1.4 × 10^−4^ in *An. melas* to ~ 1.4 × 10^−3^ in *An. arabiensis*). To approximate the ancestral estimates of *θ*, for any triple analyzed, we selected the maximum of the three corresponding estimates of *θ* as the value to pass to TASTI. The maximum was chosen to be robust to the bottleneck Fontaine et al. (2015) suspected to have occurred in the recent ancestry of some species in the complex.

Estimates of the effective population size of Anopheline species are highly variable in the literature (*e.g*., Taylor et al. 1993; Michel et al. 2006; Hodges et al. 2013). We therefore considered supplying TASTI effective population sizes across different orders of magnitude. By fitting a model with *N* ∈ {10^3^, 10^4^, 10^5^}, we found that our results were robust to effective population size. This is reasonable, since while the choice of *θ* directly affects the coalescent rate in the evolutionary process, the effective population size in essence only sets an upper bound on the migration rate. Analyses presented here assume an effective population size of *N* = 10^4^ across all populations.

We applied TASTI to the original dataset as well as 100 replicate datasets created by bootstrapping the original alignments. Parameter estimates with 95% confidence intervals for each estimated triple are shown in Table 2. Our results are concordant with earlier findings, indicating ancestral structure is frequently found among species belonging to the *Anopheles gambiae* complex. Two exceptions to this rule are between *An. coluzzii* and *An. gambiae*, and between *An. melus* and *An. merus*, as can be deduced from the markedly higher estimated ancestral migration rates indicative of lack of ancestral structure between these pairs of species under our model. The full six-species estimated species tree topology with branch confidence determined by bootstraps is presented in Figure 8. Our analysis returns the nearly fully resolved species tree ((*An. coluzzii*, *An. gambiae*), (*An. arabiensis*,(*An. melus*, *An. quadriannulatus*), *An. merus*)) with strong bootstrap support for the clades.

**Table 2:**
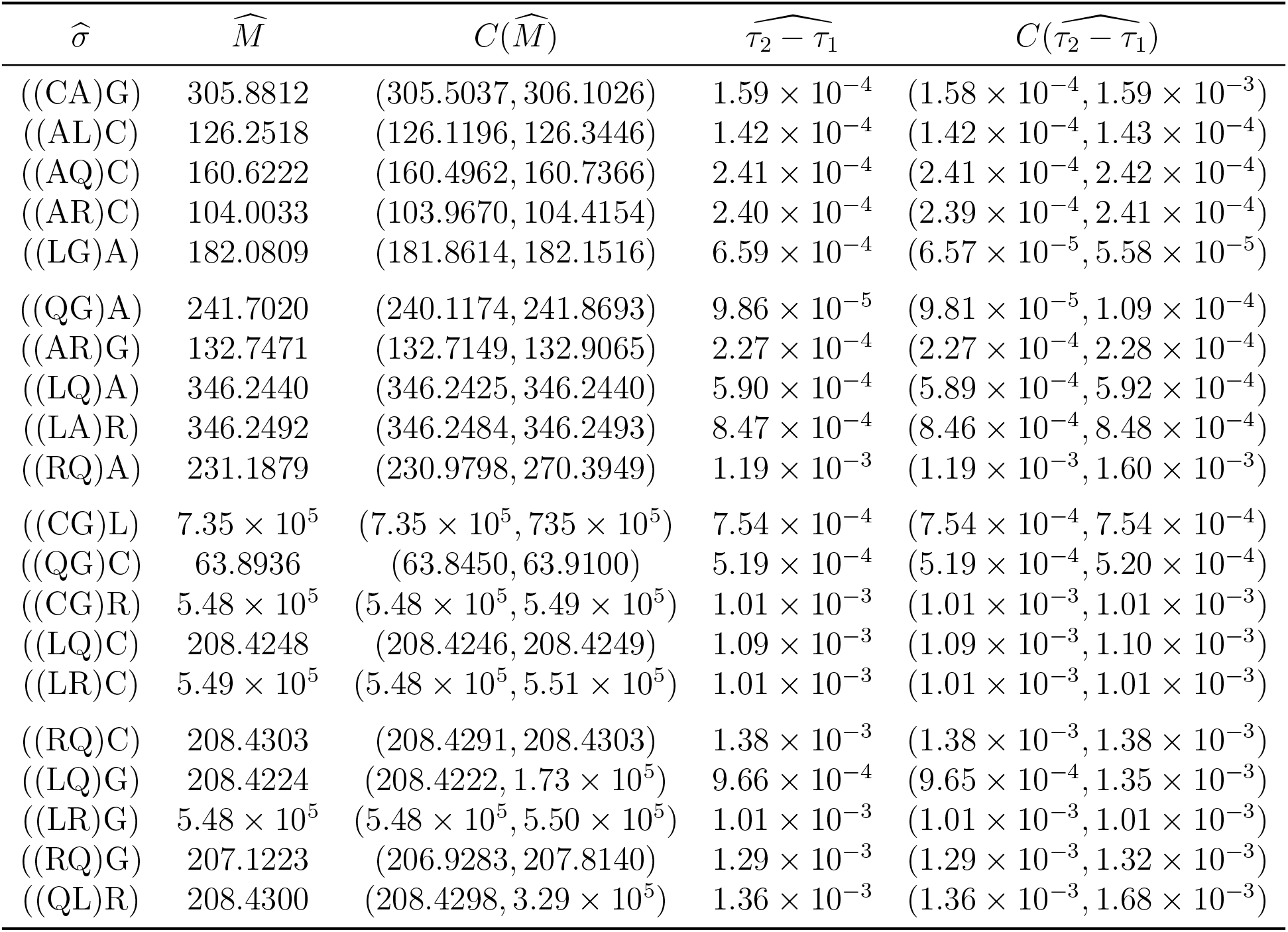
Estimated rooted species tree triples, reported in their maximum likelihood configuration, with estimates of migration rate and internal branch length are from the original data. Confidence estimates were obtained during the bootstrap analysis using 100 replicates. Species abbreviations are A (*An. arabiensis*), C (*An. coluzzii*), G (*An. gambiae*), L (*An. melus*), R (*An. merus*), and Q (*An. quadriannulatus*).

**Figure 8:**
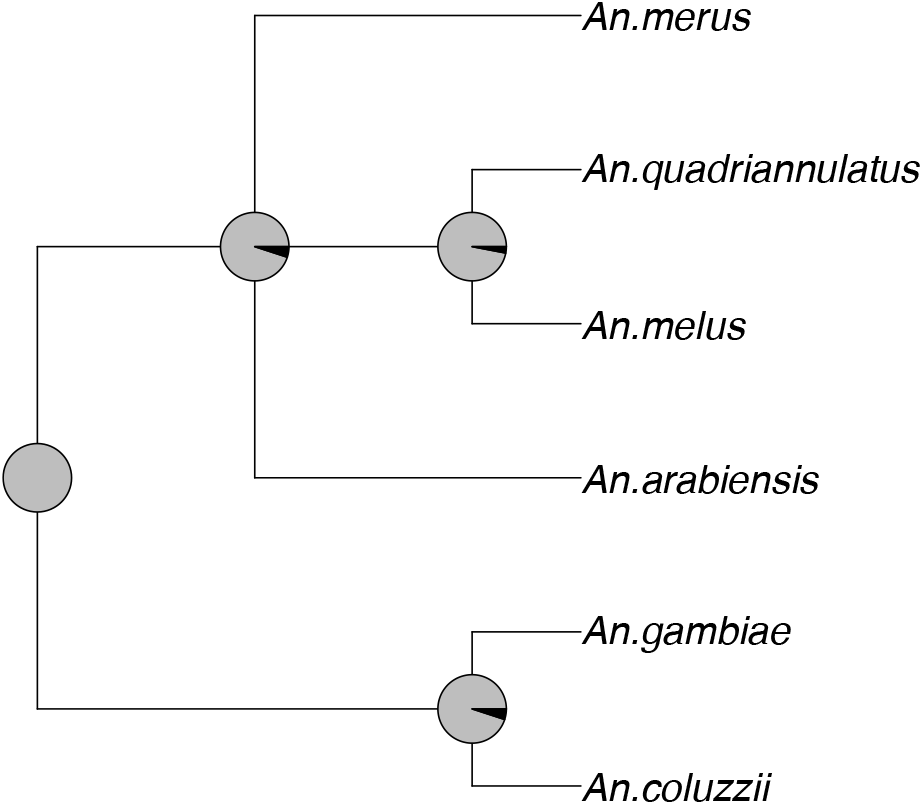
Species tree topology for the *Anopheles gambiae* complex estimated by applying the modified mincut supertree algorithm to the set of all distinct three-species tree topologies inferred by TASTI, with bootstrap branch support obtained from 100 bootstrap replicates. The proportion of gray in each pie chart corresponds to the degree of support for each clade. That is, a fully gray chart implies 100% bootstrap support for a given clade.

In line with previous work, we estimated a well-supported clade between species *An. coluzzii* and *An. gambiae*. Though our analysis and the analyses of both Fontaine et al.(2015) and Wen et al. (2016a) do not fully agree on the placement of the remaining four species in the phylogeny, because all three analyses assume a different model, the final results need not agree. It is notable, though, that the phylogeny inferred by TASTI in fact most resembles the phylogeny deduced from the X chromosome by Fontaine et al. (2015). The X chromosome of the Anopheline species does not harbor signals of introgression (Fontaine et al. 2015), so the species tree constructed from those gene trees may be more reliable in that the underlying evolutionary process is simpler to tease out. That TASTI infers a phylogeny from the autosomal data which is similar to that of the X chromosome is a reassuring result; however, identifying the correct model to assume is a non-trivial task which deserves thorough investigation in the future.

## Discussion

In this article, we developed TASTI, a scalable algorithm that can be used to infer species trees from gene trees when a group of taxa’s ancestral populations are structured.

TASTI is an improvement upon previous approaches developed for the same purpose such as GLASS, STEM, and Maximum Tree, because it achieves good performance even when gene trees are inferred. We recognize that this generalization of the standard multispecies coalescent does contain its own oversimplifications of ancestry. Specifically, the assumption of no more than two subpopulations may for some species be insufficient. Additionally, even with only two subpopulations, one could forseeably introduce more complex species tree orientations by considering other ways in which an ancestry could be structured. However, for this model, such intricacies would come at the cost of total unidentifiability of the species tree model. This issue for phylogenetic inference has been discussed at length (Chang 1996; Evans and Warnow 2004; Ho and Ané 2014; Xu and Yang 2016), and our simplified ancestral population structure model is a step in relaxing the common assumption of unstructured ancestral populations used in phylogenetics.

Notably, our extensive simulations support the idea that ancestral population structure gives rise to asymmetries in gene tree topology distributions not observed under the standard multispecies coalescent (Slatkin and Pollack 2008), but that are frequently attributed to inter-species gene flow. A substantial amount of recent research has focused on advancing models for estimating reticulate evolutionary histories caused by events such as hybridization and continuous gene flow (Huson et al. 2005; Baum 2007; Meng and Kubatko 2009; Gerard et al. 2011; Yu et al. 2014; Solís-Lemus and Ané 2016; Tian and Kubatko 2016; Wen and Nakhleh 2017; Hey et al. 2018; Long and Kubatko 2018). In contrast, though population structure is often explored in analyses of evolutionary histories (*e.g*., *F*-statistics such as *F*_*ST*_ (Wright 1949; Weir and Cockerham 1984) and software such as STRUCTURE (Pritchard et al. 2000) for exploring population structure enjoy widespread use), model-based approaches incorporating such structure within species inference frameworks have been lacking. Our analyses of the *Anopheles gambiae* complex data further demonstrate, when comparing to the work of Wen et al. (2016a) and Fontaine et al. (2015), that similar phylogenies can be inferred when assuming either a reticulate history or ancestral population structure. Indeed, the identifiability of such histories given gene tree topologies alone is severely limited or impossible. An investigation via comprehensive simulation studies of how distributions of gene tree branch lengths differ under various evolutionary assumptions such as hybridization and population structure may provide key insights into selection of an appropriate evolutionary model. Alternatively, TASTI could be reformulated in terms of a hypothesis test. That is, it may be meaningful to test for significant confidence in ancestral structure versus unstructured populations. Moreover, a different test could assess the likelihood of ancestral structure versus hybridization, though addressing such a question may be challenging based on topology data alone. Further work remains in examining how phylogenetic signals differ under these various modeling assumptions, such that an appropriate model may be determined on a case by case basis.

If possible, external information or expert opinion can be used to aid in model performance. In the case of identifiability, external knowledge can be supplied to restrict the domain of certain free parameters, namely the divergence times, such that the model becomes identifiable (Huelsenbeck et al. 2008). Models equally likely to produce a given distribution of gene tree topologies can further be evaluated by more closely examining branch lengths or molecular sequences (Yu and Nakhleh 2015). External information in the form of Bayesian prior distributions can boost the performance of gene tree inference as well (Huelsenbeck and Ronquist 2001; Huelsenbeck et al. 2004). The more accurate the input data, the closer TASTI gets to achieving optimal performance. This is especially crucial when using data on gene trees with branch lengths. Simultaneous estimation of genes trees and the species tree from multilocus data in a Bayesian hierarchical framework has also proven effective and popular (Liu and Pearl 2007; Liu 2008; Heled and Drummond 2009; Wen et al. 2016b; Zhang et al. 2017), and an extension of TASTI to such a framework may be powerful.

Some avenues are available to reduce computation time and improve performance in analyses with larger numbers of species. Maximum pseudo-likelihood methods based on groups of rooted triples have been proposed both for the standard multispecies coalescent with incomplete lineage sorting (Liu et al. 2010a) and with hybridization (Yu and Nakhleh 2015; Solís-Lemus and Ané 2016). These approaches have been shown to reduce computational burden, an important feature given that, even with greatly diminishing the computational task using a supertree approach, TASTI’s computation time grows with the number of taxa at a rate of 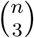. Thus, a pseudo-likelihood method for taxa with ancestral population structure would be a useful extension to our work. Finally, while we find favorable performance of the modified min cut supertree algorithm of Page (2002), the effect of alternative supertree construction algorithms (Wilkinson et al. 2005) could nevertheless be explored.

## Acknowledgements and Funding

This research was funded by National Science Foundation grant DEB-1753489, by National Institutes of Health grant R35GM128590, by the Alfred P. Sloan Foundation, by a NIGMS funded training grant on Computation, Bioinformatics and Statistics (Predoctoral Training Program T32GM102057), and by a NHGRI pre-doctoral fellowship (1F31HG010574-01). Portions of this research were conducted with Advanced CyberInfrastructure computational resources provided by the Institute for CyberScience at Pennsylvania State University.

## Availability

Independent alignments of high-depth samples from the *An. gambiae* complex, first generated by Wen et al. (2016a), are available from the Dryad Digital Repository: http://dx.doi.org/10.5061/dryad.tn47c. The VCF files for low-depth samples from the *An.gambiae* complex, detailed by Fontaine et al. (2015), are also available from the Dryad Digital Repository: http://dx.doi.org/10.5061/dryad.f4114. An R package, called TASTI, containing the functions used to implement the core analyses presented here is available for download from https://github.com/hillarykoch/TASTI.

## Notes

http://dx.doi.org/10.5061/dryad.tn47c

