## Supplementary Materials for "Maximum likelihood estimation of species trees from gene trees in the presence of ancestral population structure"

| | $a_1b_1c_1$ | $a_1b_1c_2$ | $a_1b_2c_1$ | $a_2b_1c_1$ | $a_1b_2c_2$ | $a_2b_1c_2$ | $a_2b_2c_1$ | $a_2b_2c_2$ | $(ab)_1c_1$ | $(ab)_1c_2$ |
|---|---|---|---|---|---|---|---|---|---|---|
| $a_1b_1c_1$ | $-$ | $M_{\text{ABC},2}$ | $M_{\text{ABC},2}$ | $M_{\text{ABC},2}$ | 0 | 0 | 0 | 0 | $3c_{\text{ABC},1}$ | 0 |
| $a_1b_1c_2$ | $M_{\text{ABC},1}$ | $-$ | 0 | 0 | $M_{\text{ABC},2}$ | $M_{\text{ABC},2}$ | 0 | 0 | 0 | $c_{\text{ABC},1}$ |
| $a_1b_2c_1$ | $M_{\text{ABC},1}$ | 0 | $-$ | 0 | $M_{\text{ABC},2}$ | 0 | $M_{\text{ABC},2}$ | 0 | 0 | 0 |
| $a_2b_1c_1$ | $M_{\text{ABC},1}$ | 0 | 0 | $-$ | 0 | $M_{\text{ABC},2}$ | $M_{\text{ABC},2}$ | 0 | 0 | 0 |
| $a_1b_2c_2$ | 0 | $M_{\text{ABC},1}$ | $M_{\text{ABC},1}$ | 0 | $-$ | 0 | 0 | $M_{\text{ABC},2}$ | 0 | 0 |
| $a_2b_1c_2$ | 0 | $M_{\text{ABC},1}$ | 0 | $M_{\text{ABC},1}$ | 0 | $-$ | 0 | $M_{\text{ABC},2}$ | 0 | 0 |
| $a_2b_2c_1$ | 0 | 0 | $M_{\text{ABC},1}$ | $M_{\text{ABC},1}$ | 0 | 0 | $-$ | $M_{\text{ABC},2}$ | 0 | 0 |
| $a_2b_2c_2$ | 0 | 0 | 0 | 0 | $M_{\text{ABC},2}$ | $M_{\text{ABC},1}$ | $M_{\text{ABC},1}$ | $-$ | 0 | 0 |
| $(ab)_1c_1$ | 0 | 0 | 0 | 0 | 0 | 0 | 0 | 0 | $-$ | $M_{\text{ABC},2}$ |
| $(ab)_1c_2$ | 0 | 0 | 0 | 0 | 0 | 0 | 0 | 0 | $M_{\text{ABC},1}$ | $-$ |
| $(ab)_2c_1$ | 0 | 0 | 0 | 0 | 0 | 0 | 0 | 0 | $M_{\text{ABC},1}$ | 0 |
| $(ab)_2c_2$ | 0 | 0 | 0 | 0 | 0 | 0 | 0 | 0 | 0 | $M_{\text{ABC},1}$ |
| $(ac)_1b_1$ | 0 | 0 | 0 | 0 | 0 | 0 | 0 | 0 | 0 | 0 |
| $(ac)_2b_1$ | 0 | 0 | 0 | 0 | 0 | 0 | 0 | 0 | 0 | 0 |
| $(ac)_1b_2$ | 0 | 0 | 0 | 0 | 0 | 0 | 0 | 0 | 0 | 0 |
| $(ac)_2b_2$ | 0 | 0 | 0 | 0 | 0 | 0 | 0 | 0 | 0 | 0 |
| $(bc)_1a_1$ | 0 | 0 | 0 | 0 | 0 | 0 | 0 | 0 | 0 | 0 |
| $(bc)_1a_2$ | 0 | 0 | 0 | 0 | 0 | 0 | 0 | 0 | 0 | 0 |
| $(bc)_2a_1$ | 0 | 0 | 0 | 0 | 0 | 0 | 0 | 0 | 0 | 0 |
| $(bc)_2a_2$ | 0 | 0 | 0 | 0 | 0 | 0 | 0 | 0 | 0 | 0 |

| | $(ab)_2c_1$ | $(ab)_2c_2$ | $(ac)_1b_1$ | $(ac)_1b_2$ | $(ac)_2b_1$ | $(ac)_2b_2$ | $(bc)_1a_1$ | $(bc)_1a_2$ | $(bc)_2a_1$ | $(bc)_2a_2$ |
|---|---|---|---|---|---|---|---|---|---|---|
| $a_1b_1c_1$ | 0 | 0 | $3c_{\text{ABC},1}$ | 0 | 0 | 0 | $3c_{\text{ABC},1}$ | 0 | 0 | 0 |
| $a_1b_1c_2$ | 0 | 0 | 0 | 0 | 0 | 0 | 0 | 0 | 0 | 0 |
| $a_1b_2c_1$ | 0 | 0 | 0 | $c_{\text{ABC},1}$ | 0 | 0 | 0 | 0 | 0 | 0 |
| $a_2b_1c_1$ | 0 | 0 | 0 | 0 | 0 | 0 | 0 | $c_{\text{ABC},1}$ | 0 | 0 |
| $a_1b_2c_2$ | 0 | 0 | 0 | 0 | 0 | 0 | 0 | 0 | $c_{\text{ABC},2}$ | 0 |
| $a_2b_1c_2$ | 0 | 0 | 0 | 0 | $c_{\text{ABC},2}$ | 0 | 0 | 0 | 0 | 0 |
| $a_2b_2c_1$ | $c_{\text{ABC},2}$ | 0 | 0 | 0 | 0 | 0 | 0 | 0 | 0 | 0 |
| $a_2b_2c_2$ | 0 | $3c_{\text{ABC},2}$ | 0 | 0 | 0 | $3c_{\text{ABC},2}$ | 0 | 0 | 0 | $3c_{\text{ABC},2}$ |
| $(ab)_1c_1$ | $M_{\text{ABC},2}$ | 0 | 0 | 0 | 0 | 0 | 0 | 0 | 0 | 0 |
| $(ab)_1c_2$ | 0 | $M_{\text{ABC},2}$ | 0 | 0 | 0 | 0 | 0 | 0 | 0 | 0 |
| $(ab)_2c_1$ | $-$ | $M_{\text{ABC},2}$ | 0 | 0 | 0 | 0 | 0 | 0 | 0 | 0 |
| $(ab)_2c_2$ | $M_{\text{ABC},1}$ | $-$ | 0 | 0 | 0 | 0 | 0 | 0 | 0 | 0 |
| $(ac)_1b_1$ | 0 | 0 | $-$ | $M_{\text{ABC},2}$ | $M_{\text{ABC},2}$ | 0 | 0 | 0 | 0 | 0 |
| $(ac)_1b_2$ | 0 | 0 | $M_{\text{ABC},1}$ | $-$ | 0 | $M_{\text{ABC},2}$ | 0 | 0 | 0 | 0 |
| $(ac)_2b_1$ | 0 | 0 | $M_{\text{ABC},1}$ | 0 | $-$ | $M_{\text{ABC},2}$ | 0 | 0 | 0 | 0 |
| $(ac)_2b_2$ | 0 | 0 | 0 | $M_{\text{ABC},1}$ | $M_{\text{ABC},1}$ | $-$ | 0 | 0 | 0 | 0 |
| $(bc)_1a_1$ | 0 | 0 | 0 | 0 | 0 | 0 | $-$ | $M_{\text{ABC},2}$ | $M_{\text{ABC},2}$ | 0 |
| $(bc)_1a_2$ | 0 | 0 | 0 | 0 | 0 | 0 | $M_{\text{ABC},1}$ | $-$ | 0 | $M_{\text{ABC},2}$ |
| $(bc)_2a_1$ | 0 | 0 | 0 | 0 | 0 | 0 | $M_{\text{ABC},1}$ | 0 | $-$ | $M_{\text{ABC},2}$ |
| $(bc)_2a_2$ | 0 | 0 | 0 | 0 | 0 | 0 | 0 | $M_{\text{ABC},1}$ | $M_{\text{ABC},1}$ | $-$ |

Figure S1: Instantaneous rate matrix $\mathbf{Q}_{\text{ABC}}$ describing the time to the first coalescence above the root, if none happen along the internal branch.

Table S1: Median parameter estimates of 100 simulated replicates from $K \in \{100,\ 500,\ 1000\}$ loci across all migration settings.

| | | Exact topologies | | Inferred topologies | | Exact gene trees | | Inferred gene trees | |
|---|---|---|---|---|---|---|---|---|---|
| $M$ | loci | $\widehat{M}$ | $\widehat{\tau_2 - \tau_1}$ | $\widehat{M}$ | $\widehat{\tau_2 - \tau_1}$ | $\widehat{M}$ | $\widehat{\tau_2 - \tau_1}$ | $\widehat{M}$ | $\widehat{\tau_2 - \tau_1}$ |
| 0.5 | 100 | $9.12\times10^{-4}$ | $9.12\times10^{-4}$ | 1.15 | $9.12\times10^{-4}$ | 0.505 | $2.76\times10^{-3}$ | 0.485 | $1.50\times10^{-3}$ |
| | 500 | 0.282 | $9.12\times10^{-4}$ | 0.751 | $2.78\times10^{-3}$ | 0.500 | $2.75\times10^{-3}$ | 0.482 | $1.00\times10^{-3}$ |
| | 1000 | 0.258 | $9.66\times10^{-4}$ | 0.565 | $2.79\times10^{-3}$ | 0.499 | $2.75\times10^{-3}$ | 0.495 | $1.00\times10^{-3}$ |
| 5 | 100 | 2.65 | $9.12\times10^{-4}$ | 5.24 | $9.12\times10^{-4}$ | 4.92 | $2.77\times10^{-3}$ | 4.87 | $1.51\times10^{-3}$ |
| | 500 | 3.78 | $9.12\times10^{-4}$ | 5.11 | $9.12\times10^{-4}$ | 4.93 | $2.75\times10^{-3}$ | 4.83 | $1.00\times10^{-3}$ |
| | 1000 | 4.31 | $9.12\times10^{-4}$ | 4.97 | $9.12\times10^{-4}$ | 4.99 | $2.75\times10^{-3}$ | 4.87 | $1.00\times10^{-3}$ |
| 50 | 100 | 36.7 | $9.12\times10^{-4}$ | 45.9 | $9.12\times10^{-4}$ | 43.8 | $2.76\times10^{-3}$ | 39.9 | $1.72\times10^{-3}$ |
| | 500 | 39.5 | $9.12\times10^{-4}$ | 45.4 | $9.12\times10^{-4}$ | 43.7 | $2.75\times10^{-3}$ | 39.4 | $1.50\times10^{-3}$ |
| | 1000 | 39.8 | $9.12\times10^{-4}$ | 46.2 | $9.12\times10^{-4}$ | 44.1 | $2.75\times10^{-3}$ | 40.0 | $1.00\times10^{-3}$ |
| $4N/\theta$ | 100 | $1.08\times10^{3}$ | $9.12\times10^{-4}$ | $8.85\times10^{2}$ | $9.12\times10^{-4}$ | $1.50\times10^{4}$ | $2.74\times10^{-3}$ | $1.05\times10^{3}$ | $2.16\times10^{-3}$ |
| | 500 | $9.42\times10^{2}$ | $9.12\times10^{-4}$ | $9.02\times10^{2}$ | $9.12\times10^{-4}$ | $1.55\times10^{4}$ | $2.73\times10^{-3}$ | 660 | $1.77\times https://www.overleaf.com/21461485kzkqsqrwkcyw10^{-3}$ |
| | 1000 | $9.64\times10^{2}$ | $9.12\times10^{-4}$ | $8.90\times10^{2}$ | $9.12\times10^{-4}$ | $5.40\times10^{4}$ | $2.74\times10^{-3}$ | 657 | $1.79\times10^{-3}$ |

Table S2: 2.5% and 97.5% quantiles for 100 simulated replicates from $K \in \{100,\ 500,\ 1000\}$ loci across all migration settings.

| | | Exact topologies | | Inferred topologies | | Exact gene trees | | Inferred gene trees | |
|---|---|---|---|---|---|---|---|---|---|
| $M$ | loci | $\widehat{M}$ | $\widehat{\tau_2 - \tau_1}$ | $\widehat{M}$ | $\widehat{\tau_2 - \tau_1}$ | $\widehat{M}$ | $\widehat{\tau_2 - \tau_1}$ | $\widehat{M}$ | $\widehat{\tau_2 - \tau_1}$ |
| 0.5 | 100 | $(9.12\times10^{-4}, 2.77)$ | $(9.12\times10^{-4}, 6.48\times10^{-3})$ | $(9.12\times10^{-4}, 3.1315738\times10^{7})$ | $(9.12\times10^{-4}, 6.48\times10^{-2})$ | (0.433, 0.606) | $(4.00\times10^{-5}, 1.56\times10^{-3})$ | (0.269, 0.659) | $(2.91\times10^{-4}, 2.33\times10^{-3})$ |
| | 500 | $(9.12\times10^{-4}, 1.04)$ | $(9.12\times10^{-4}, 6.48\times10^{-3})$ | $(0.106, 2.50\times10^{7})$ | $(9.12\times10^{-4}, 6.48\times10^{-3})$ | (0.466, 0.536) | $(3.95\times10^{-5}, 1.54\times10^{-3})$ | (0.333, 0.549) | $(5.16\times10^{-4}, 1.16\times10^{-3})$ |
| | 1000 | (0.0531, 0.922) | $(9.12\times10^{-4}, 6.48\times1-^{-3})$ | $(0.185, 3.49\times10^{7})$ | $(9.12\times10^{-4}, 6.48\times10^{-3})$ | (0.458, 0.529) | $(8.40\times10^{-6}, 3.22\times10^{-4})$ | (0.370, 0.539) | $(5.29\times10^{-4}, 1.65\times10^{-3})$ |
| 5 | 100 | $(9.12\times10^{-4}, 9.00)$ | $(9.12\times10^{-4}, 6.48\times10^{-3})$ | $(0.792, 3.45\times10^{7})$ | $(9.12\times10^{-4}, 6.48\times10^{-3})$ | (4.22, 5.97) | $(6.60\times10^{-5}, 2.57\times10^{-3})$ | (4.02, 6.46) | $(6.40\times10^{-5}, 2.49\times10^{-3})$ |
| | 500 | (1.78, 6.47) | $(9.12\times10^{-4}, 4.30\times10^{-3})$ | $(1.77, 2.26\times10^{7})$ | $(9.12\times10^{-4}, 5.80\times10^{-3})$ | (4.49, 5.38) | $(1.59\times10^{-3}, 2.72\times10^{-3})$ | (4.34, 5.42) | $(2.70\times10^{-4}, 1.51\times10^{-3})$ |
| | 1000 | (2.36, 5.78) | $(9.12\times10^{-4}, 3.21\times10^{-3})$ | $(3.07, 2.72\times10^{7})$ | $(9.12\times10^{-4}, 2.95\times10^{-3})$ | (4.65, 5.30) | $(2.85\times10^{-5}, 1.11\times10^{-3})$ | (4.43, 5.31) | $(2.70\times10^{-4}, 1.51\times10^{-3})$ |
| 50 | 100 | (22.8, 57.8) | $(9.12\times10^{-4}, 3.23\times10^{-3})$ | $(23.6, 2.93\times10^{7})$ | $(9.12\times10^{-4}, 3.73\times10^{-3})$ | (35.8, 55.1) | $(4.80\times10^{-6}, 1.86\times10^{-4})$ | (29.3, 59.7) | $(5.73\times10^{-4}, 3.33\times10^{-3})$ |
| | 500 | (30.7, 48.3) | $(9.12\times10^{-4}, 1.65\times10^{-3})$ | $(32.6, 2.89\times10^{7})$ | $(9.12\times10^{-4}, 1.78\times10^{-3})$ | (36.4, 47.9) | $(3.34\times10^{-5}, 1.30\times10^{-3})$ | (34.0, 50.0) | $(5.25\times10^{-4}, 1.49\times10^{-3})$ |
| | 1000 | (33.8, 46.8) | $(9.12\times10^{-4}, 1.06\times10^{-3})$ | $(37.8, 3.22\times10^{7})$ | $(9.12\times10^{-4}, 1.44\times10^{-3})$ | (35.4, 47.9) | $(2.02\times10^{-4}, 1.27\times10^{-3})$ | (35.8, 45.0) | $(4.78\times10^{-5}, 1.86\times10^{-3})$ |
| $4N/\theta$ | 100 | $(181, 2.08\times10^{7})$ | $(9.12\times10^{-4}, 2.37\times10^{-3})$ | $(176, 2.58\times10^{7})$ | $(9.12\times10^{-4}, 2.44\times10^{-3})$ | $(225, 10^{7})$ | $(3.43\times10^{-4}, 1.17\times10^{-3})$ | $(214, 1.52\times10^{4})$ | $(1.83\times10^{-5}, 7.14\times10^{-4})$ |
| | 500 | $(443, 2.36\times10^{7})$ | $(9.12\times10^{-4}, 1.34\times10^{-3})$ | $(364, 2.04\times10^{7})$ | $(9.12\times10^{-4}, 1.44\times10^{-3})$ | $(238, 10^{7})$ | $(3.19\times10^{-4}, 1.22\times10^{-3})$ | $(236, 1.23\times10^{3})$ | $(1.53\times10^{-5}, 5.94\times10^{-4})$ |
| | 1000 | $(574, 2.62\times10^{7})$ | $(9.12\times10^{-4}, 1.05\times10^{-3})$ | $(452, 2.39\times10^{7})$ | $(9.12\times10^{-4}, 1.02\times10^{-3})$ | $(243, 10^{7})$ | $(3.26\times10^{-4}, 2.45\times10^{-3})$ | $(230, 1.38\times10^{3})$ | $(1.34\times10^{-5}, 5.23\times10^{-4})$ |

### Parameter estimation for gene trees with branch lengths

Numerical optimization in the parameter space under the setting where branch lengths are included in the model is challenging. Indeed, in our simulations, when only gene tree topologies were used, there were just two parameters to optimize over (the internal branch length and the symmetric migration rate between subpopulations). However, with the introduction of branch lengths comes a third parameter, because we now need to infer divergence times $\tau_1$ and $\tau_2$ separately, rather than only the internal branch length $\tau_2 - \tau_1$. While direct numerical optimization is a reasonable approach, we observed that the estimates of $\tau_1$ and $\tau_2$ were sometimes inaccurate when this technique was employed. In particular, the estimate of $\tau_1$ was frequently pushed to zero while $\tau_2$ was pushed toward its upper bound, though these parameters do not necessarily maximize the likelihood.

As an alternative, we considered optimization based on a three-dimensional grid search throughout the parameter space. Of course, the finer the grid, the more accurate the parameter estimates will be, but at the cost of greater computational burden. Following recommendations based on simulations from Mailund et al. (2012), we coarsened the migration parameter space into 50 bins, and coarsened the divergence times parameter space into 20 bins each. This procedure frequently produced more stable and accurate parameter estimates that those presented in the main body of our paper. Here, we present simulation results based on performance using grid search optimization in Figures S2 (column 3) and S3, and median parameter estimates and 95% confidence intervals in Tables S3 and S4, respectively. Interestingly, although parameter estimates are less stable under numerical optimization when compared to the three-dimensional grid search, the final species tree inference is often better using this technique.

Table S3: Median parameter estimates of 100 simulated replicates from $K \in \{100, 500, 1000\}$ loci across all migration settings obtained using grid search optimization.

| | | Exact gene trees | | Inferred gene trees | |
|---|---|---|---|---|---|
| $M$ | loci | $\widehat{M}$ | $\widehat{\tau_2 - \tau_1}$ | $\widehat{M}$ | $\widehat{\tau_2 - \tau_1}$ |
| 0.5 | 100 | 0.60 | 0 | 0.60 | 0 |
| | 500 | 0.60 | 0 | 0.60 | 0 |
| | 1000 | 0.60 | 0 | 0.60 | 0 |
| 5 | 100 | 3.01 | 0 | 3.01 | 0 |
| | 500 | 3.01 | $4.42\times10^{-4}$ | 3.01 | 0 |
| | 1000 | 3.01 | $4.47\times10^{-4}$ | 3.01 | 0 |
| 50 | 100 | 75.5 | 0 | 75.5 | 0 |
| | 500 | 75.5 | 0 | 75.5 | 0 |
| | 1000 | 75.5 | 0 | 75.5 | 0 |
| $4N/\theta$ | 100 | $1.20\times10^{6}$ | $2.82\times10^{-4}$ | $1.90\times10^{3}$ | $1.99\times10^{-4}$ |
| | 500 | $1.20\times10^{6}$ | $2.78\times10^{-4}$ | $3.78\times10^{2}$ | $1.23\times10^{-4}$ |
| | 1000 | $1.20\times10^{6}$ | $2.76\times10^{-4}$ | $3.78\times10^{2}$ | $2.19\times10^{-4}$ |

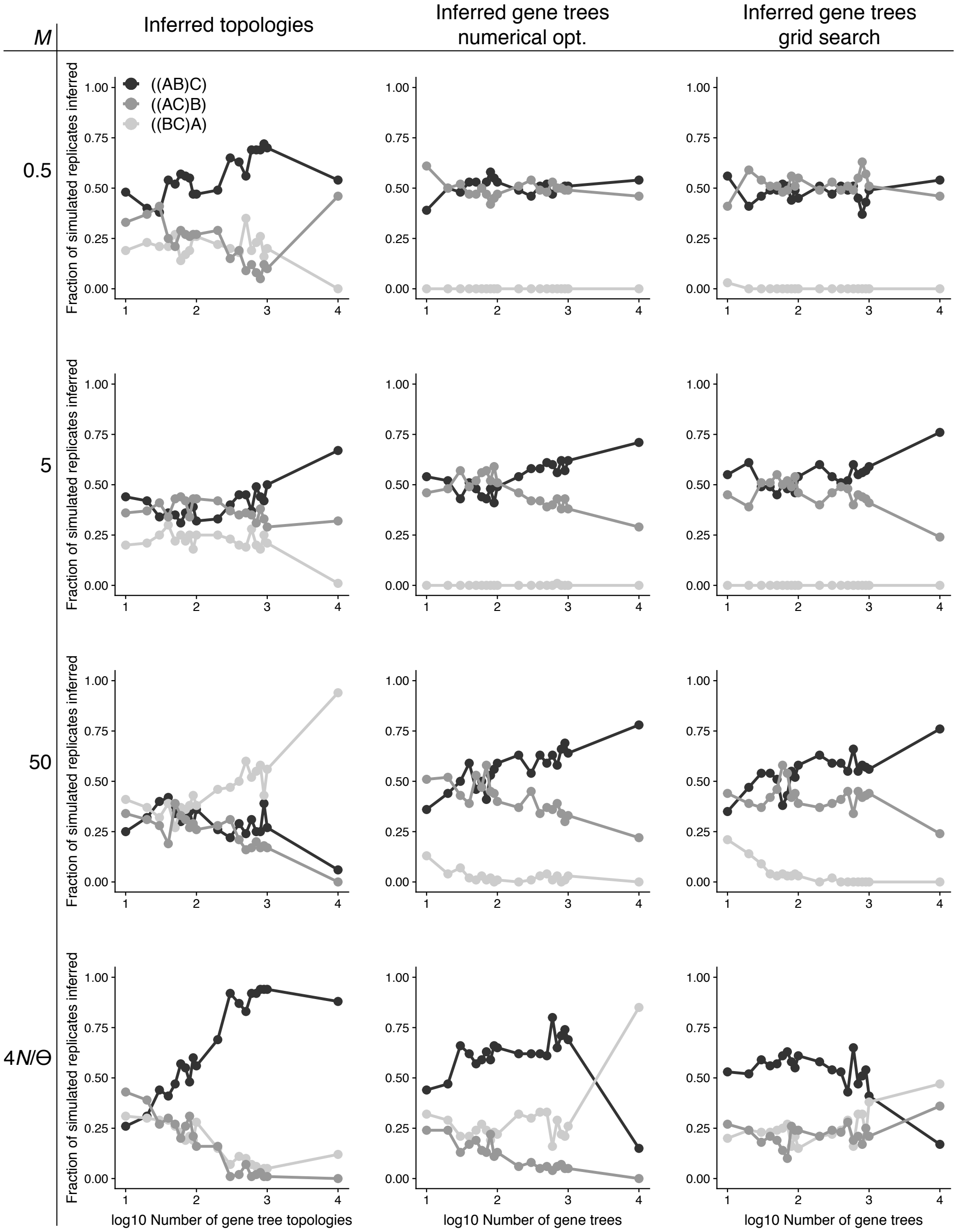


Figure S2: Accuracy of the maximum likelihood estimator of $\sigma$ as a function of the number of input gene trees. Accuracy is based on the proportion of 100 simulated replicates of $K$ loci (before filtering "star trees"), with $K$ ranging from 10 to $10^4$, where our method inferred a specific species tree topology under a scenario with $\tau_1 = 2.5 \times 10^{-3}$ and $\tau_2 = 2.75 \times 10^{-3}$. Unlike the results displayed in Figures 4 and S3, gene trees were inferred from 0.5 kb regions.

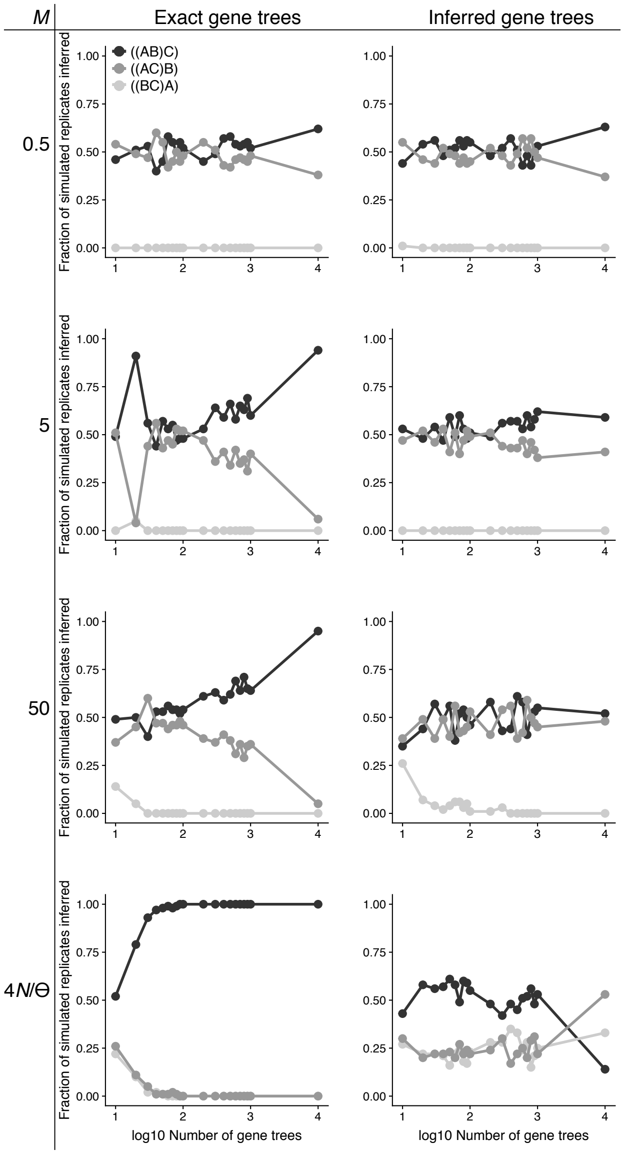


Figure S3: Accuracy of the maximum likelihood estimator of $\sigma$ as a function of the number of input gene trees. Accuracy is based on the proportion of 100 simulated replicates of $K$ loci (before filtering "star trees"), with $K$ ranging from 10 to $10^4$, where our method inferred a specific species tree topology under a scenario with $\tau_1 = 2.5 \times 10^{-3}$ and $\tau_2 = 2.75 \times 10^{-3}$. Unlike the results displayed in Figure 4, species trees were obtained using a grid search as opposed to a numerical optimization procedure.

Table S4: 2.5% and 97.5% quantiles for 100 simulated replicates from $K \in \{100, 500, 1000\}$ loci across all migration settings obtained using grid search optimization.

| | | Exact gene trees | | Inferred gene trees | |
|---|---|---|---|---|---|
| $M$ | loci | $C(\widehat{M})$ | $C(\widehat{\tau_2 - \tau_1})$ | $C(\widehat{M})$ | $C(\widehat{\tau_2 - \tau_1})$ |
| 0.5 | 100 | (0.600, 0.600) | (0, $2.45\times10^{-3}$) | (0.600, 0.600) | (0, $1.23\times10^{-3}$) |
| | 500 | (0.600, 0.600) | (0, $2.45\times10^{-3}$) | (0.600, 0.600) | (0, $9.89\times10^{-4}$) |
| | 1000 | (0.600, 0.600) | (0, $1.70\times10^{-3}$) | (0.600, 0.600) | (0, $4.16\times10^{-4}$) |
| 5 | 100 | (3.01, 3.01) | (0, $2.65\times10^{-3}$) | (3.01, 3.01) | (0, $2.57\times10^{-3}$) |
| | 500 | (3.01, 3.01) | (0,$2.59\times10^{-3}$) | (3.01, 3.01) | (0, $1.37\times10^{-3}$) |
| | 1000 | (3.01, 3.01) | (0, $2.63\times10^{-3}$) | (3.01, 3.01) | (0, $1.45\times10^{-3}$) |
| 50 | 100 | (75.5, 75.5) | (0, $2.64\times10^{-4}$) | (15.1, 75.5) | (0, $1.74\times10^{-3}$) |
| | 500 | (75.5, 75.5) | (0, $3.21\times10^{-4}$) | (75.5, 75.5) | (0, $8.71\times10^{-4}$) |
| | 1000 | (75.5, 75.5) | (0, $2.37\times10^{-4}$) | (75.5, 75.5) | (0, $1.02\times10^{-3}$) |
| $4N/\theta$ | 100 | ($9.51\times10^{3}$, $6.00\times10^{7}$) | ($1.42\times10^{-4}$, $4.25\times10^{-4}$) | (378, $951\times10^{3}$) | (0, $1.22\times10^{-3}$) |
| | 500 | ($9.51\times10^{3}$, $6.00\times10^{7}$) | ($2.06\times10^{-4}$, $2.86\times10^{-4}$) | (378, $1.90\times10^{3}$) | (0, $7.77\times10^{-4}$) |
| | 1000 | ($9.51\times10^{3}$, $6.00\times10^{7}$) | ($2.75\times10^{-4}$, $2.81\times10^{-4}$) | (378, $1.90\times10^{3}$) | (0, $6.05\times10^{-4}$) |

### Bounding the length of the internal branch of the species tree in the low signal setting

When using exact gene tree topologies for input data, and when migration is at its lowest ($M = 0.5$), TASTI never infers the species tree topology ((BC)A) as the species tree topology (Fig. 4, first column). One may be surprised that in this setting we do not infer this species tree topology, as $((bc)a)$ is by far the most prevalent sampled gene tree topology (indeed, for small numbers of loci $K$, it is often the only gene tree topology). However, by declaring an upper bound on the internal branch length as described in the *Implementation* section, we guaranteed that we must observe some amount of gene tree discordance in our

Table S5: Accuracy of the maximum likelihood estimator as a function of migration rate $M = 4Nm/\theta$. Results were obtained using exact gene trees and gene trees inferred from 1 kb sequences and a grid search optimization procedure. Accuracy is based on the percentage of 100 simulated replicates of $10^3$ loci that inferred a specific species tree topology under a scenario with $\tau_1 = 2.5 \times 10^{-3}$ and $\tau_2 = 5.25 \times 10^{-3}$.

| | Exact gene trees | | | Inferred gene trees | | |
|---|---|---|---|---|---|---|
| $M$ | ((AB)C) | ((BC)A) | ((AC)B) | ((AB)C) | ((BC)A) | ((AC)B) |
| 0.5 | 94 | 0 | 6 | 78 | 0 | 22 |
| 5 | 100 | 0 | 0 | 97 | 0 | 3 |
| 50 | 100 | 0 | 0 | 99 | 0 | 1 |
| $4N/\theta$ | 100 | 0 | 0 | 83 | 11 | 6 |

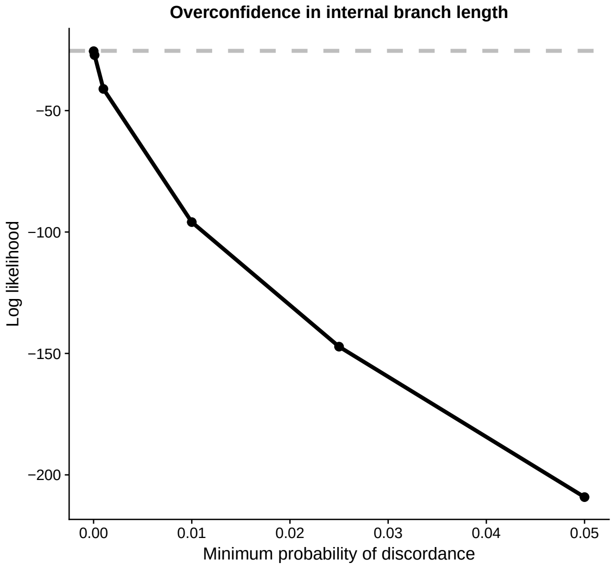


Figure S4: Examination of the effect of minimum probability of gene tree discordance on species tree inference when migration between subpopulations is very low. The mean log likelihood of the truth across 100 simulated replicates of $10^3$ loci known with certainty when $M = 0.5$ and $\tau_2 - \tau_1 = 2.5 \times 10^{-4}$ is plotted in a dashed line against the log likelihood of an alternative species tree where the ancestral species are unstructured and taxa B and C are sister species. As the minimum probability of discordance goes to 0 (implying a longer internal branch length), the likelihood of the incorrect species tree ((BC)A) exceeds the likelihood of the true tree ((AB)C).

sample. The distribution of gene tree topologies when $M = 0.5$, though, is almost never discordant with species tree ((BC)A), and thus we are left to choose between the remaining two topologies. However, if the signal is too low, then TASTI cannot distinguish between species tree topologies ((AB)C) and ((AC)B). In short, overconfidence implied by a lower bound on the probability of gene tree discordance essentially results in an almost equal chance of inferring either the true species tree topology ((AB)C), or the incorrect species tree topology ((AC)B).

We tested this claim by simulating 100 replicates of $10^3$ exact gene trees under our model with $M = 0.5$, but let the probability of discordance go to zero, corresponding to an upper bound on the internal branch length going to infinity. Specifically, we computed the mean log likelihood of the truth ($\tau_2 - \tau_1 = 2.5 \times 10^{-4}$, $M = 0.5$) across all replicates. Then, we computed the upper bound on the length of the internal branch for minimum probabilities of discordance $\mathbb{P}[G \neq \sigma] \in \{10^{-5}, 10^{-4}, 10^{-3}, 10^{-2}, 0.025, 0.05\}$. We calculated the log likelihood using a migration rate of $M = 4N/\theta$ and an internal branch length corresponding to the upper bound evaluated under each of these scenarios. Results from this simulation supported our claim (Fig. S4); if we do not assume some minimal amount of discordance, then TASTI can infer the species tree topology ((BC)A) under the scenario illustrated in Figure 1.

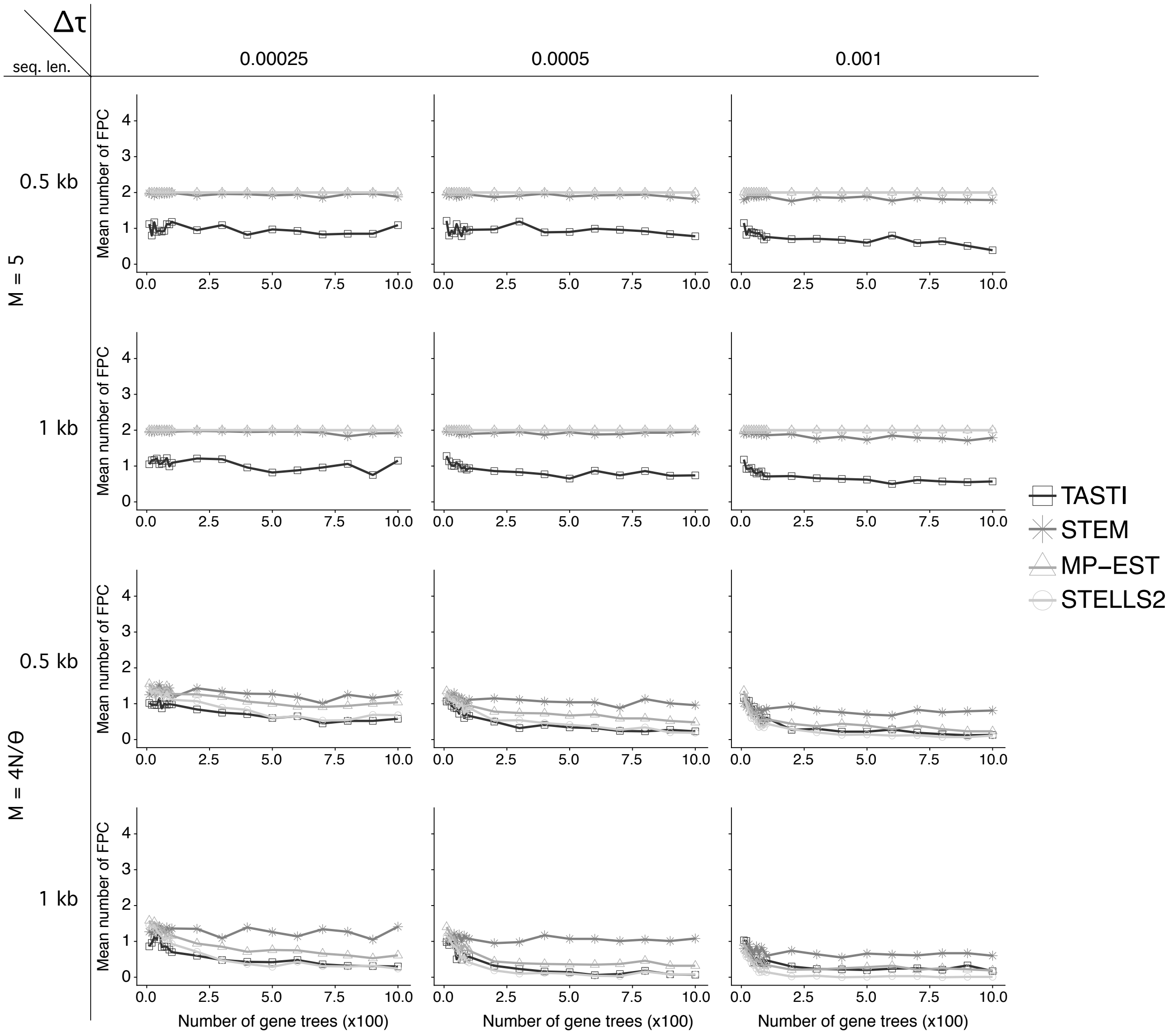


Figure S5: Mean number of false positive clades (FPC) computed between the inferred species tree topology and the true species tree topology across all supertree simulation settings by TASTI, MP-EST, STELLS2, and STEM2.0.

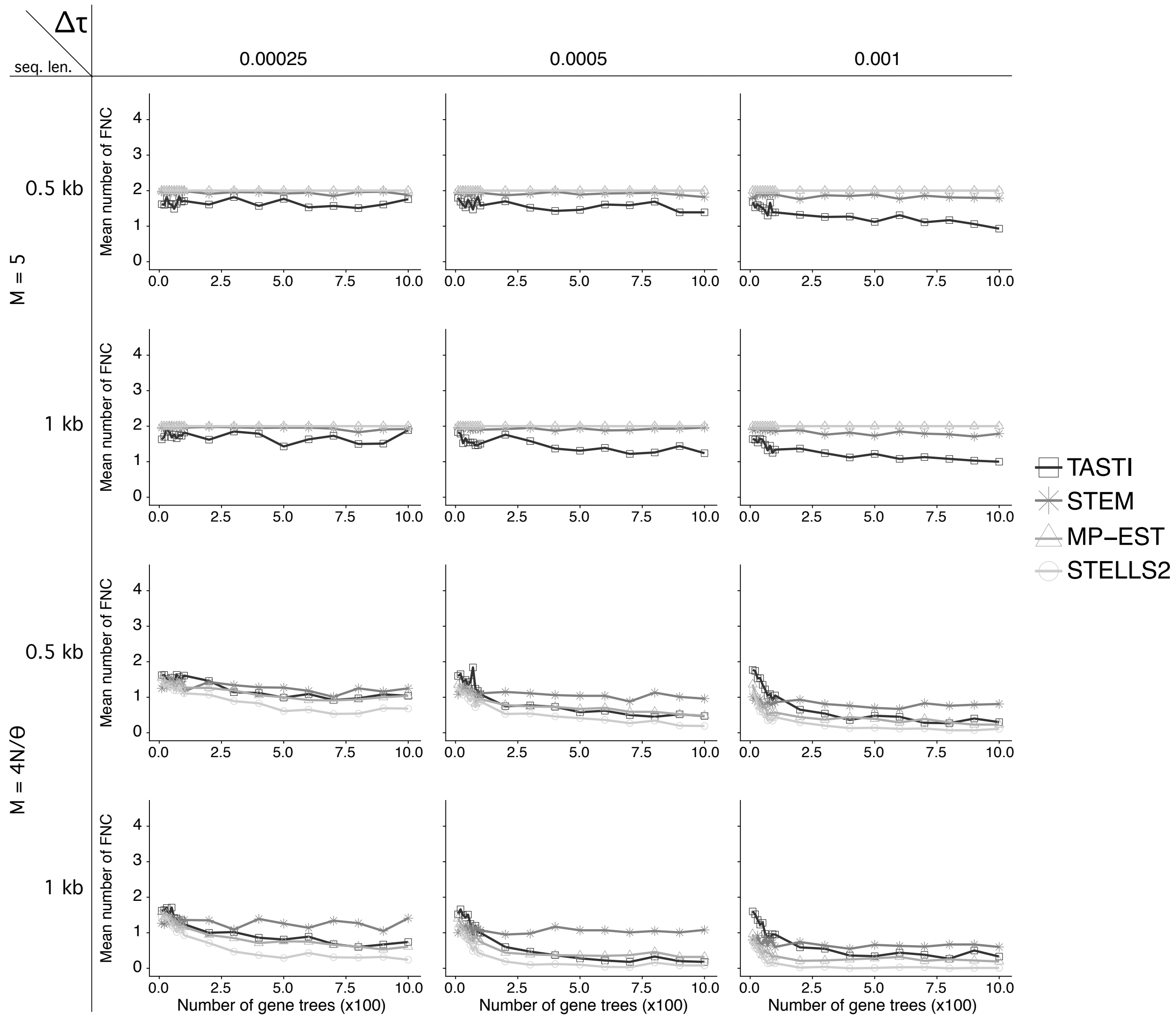


Figure S6: Mean number of false negative clades (FNC) computed between the inferred species tree topology and the true species tree topology across all supertree simulation settings by TASTI, MP-EST, STELLS2, and STEM2.0.

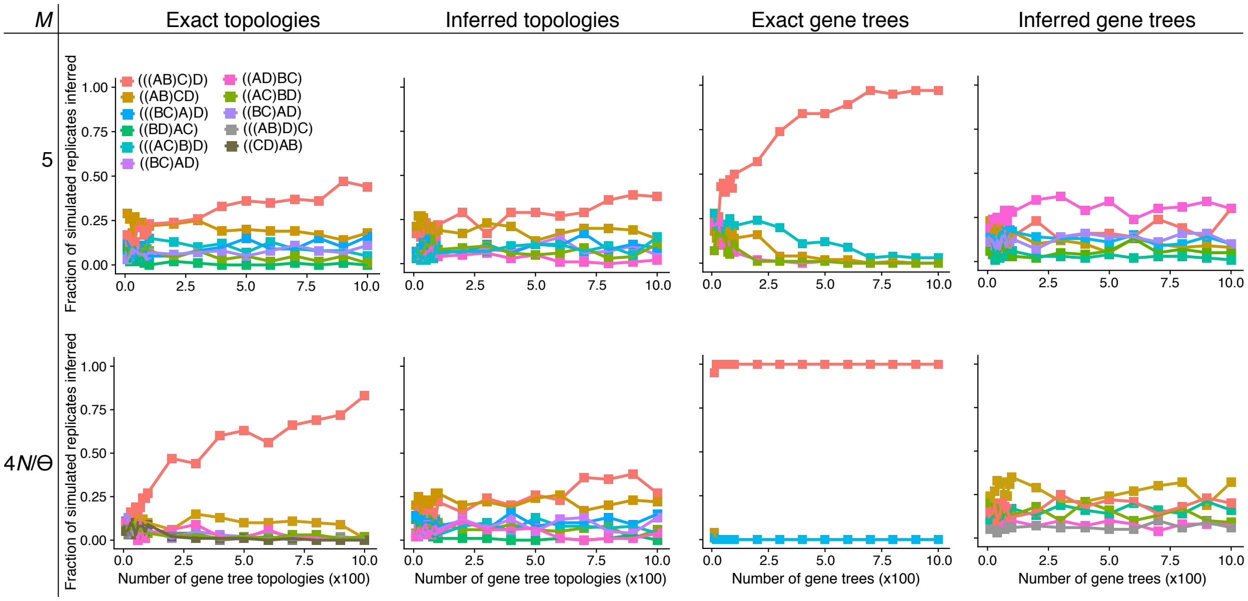


Figure S7: Accuracy of the maximum likelihood estimator of $\sigma$ as a function of the number of input gene trees. Accuracy is based on the proportion of 100 simulated replicated of $K$ loci, with $K$ ranging from 10 to $10^3$. We considered a migration rate $M = 5$ as well as the unstructured scenario. For plotting purposes, for a given simulation setting, if a large number of different topologies were inferred across the entire simulation, we only plotted topologies which occurred at least 10% of the time for a given set of 100 replicates and given number of simulated loci.

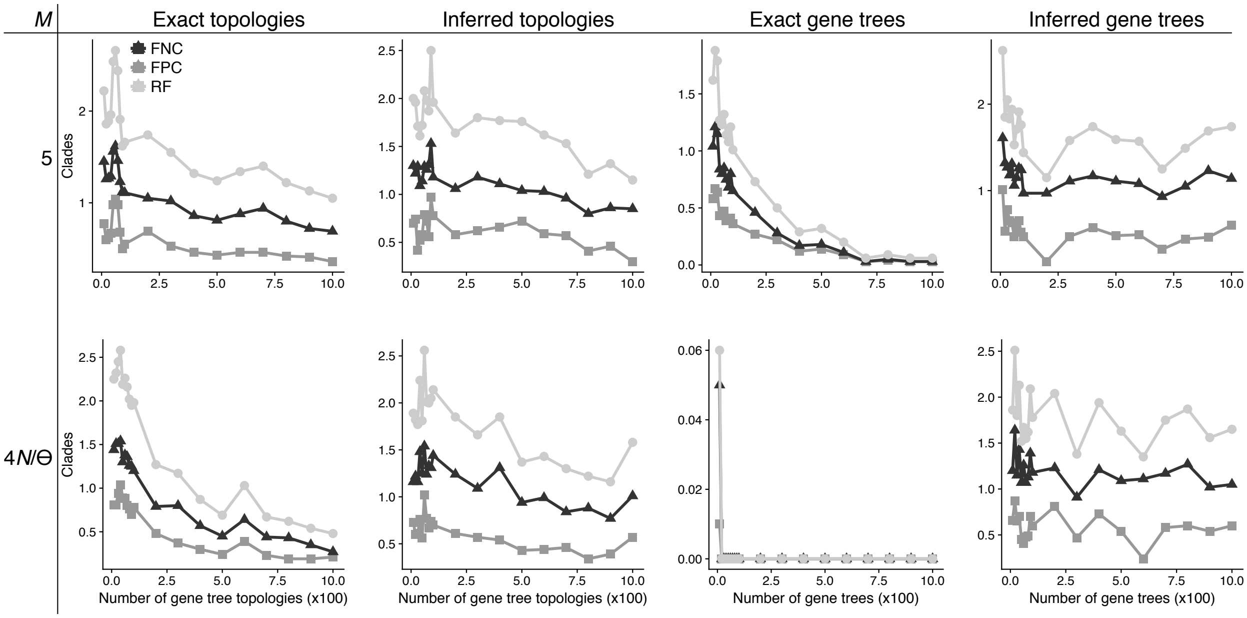


Figure S8: Mean Robinson-Foulds (RF) distance, mean number of false positive clades (FPC), and mean number of false negative clades (FNC) computed between the inferred species tree topology and the true species tree topology across all supertree simulation settings.

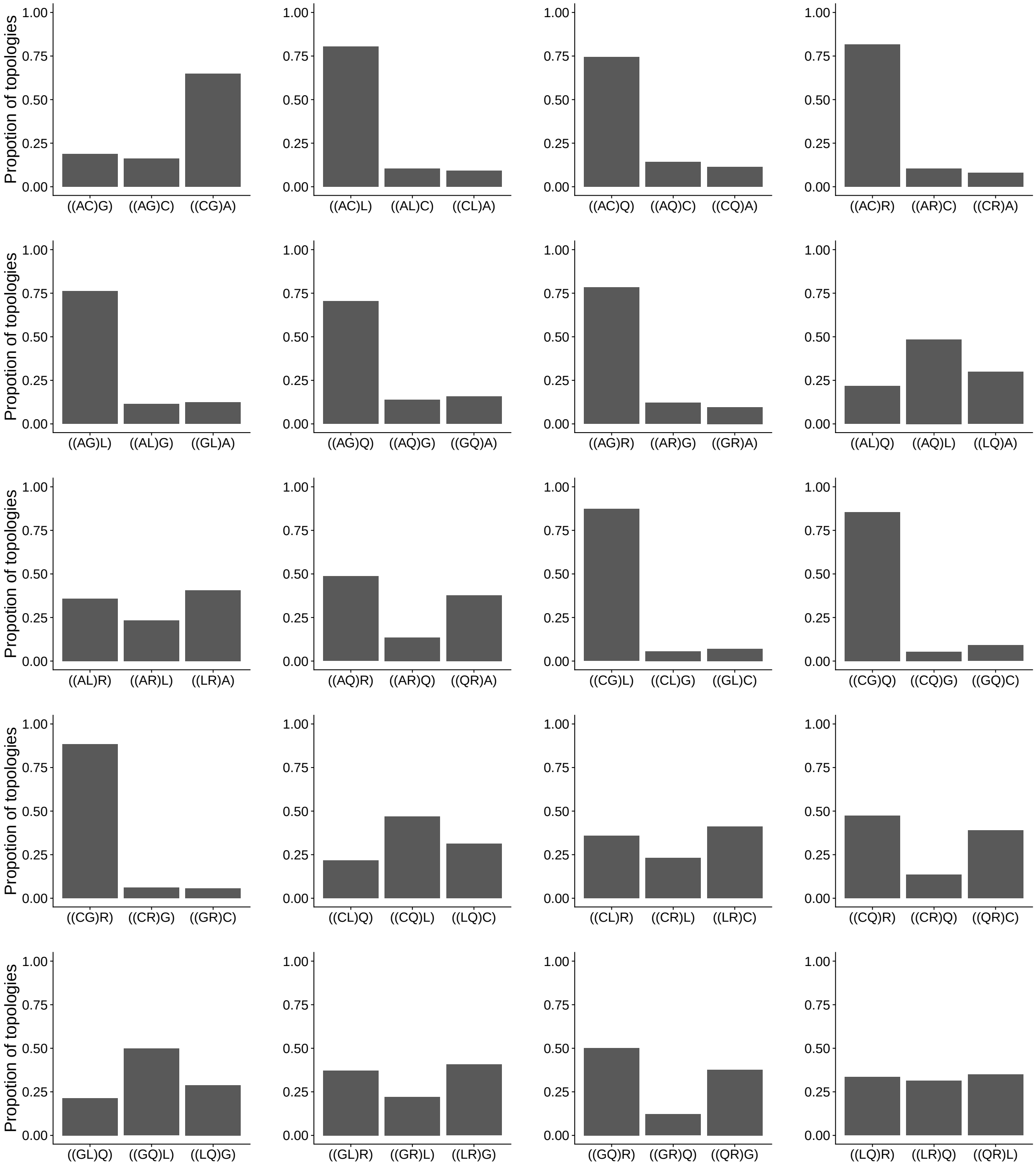


Figure S9: Distribution of gene tree topology triples for *Anopheles* data. Gene trees were estimated from genome-wide data and their constituent gene tree triples were extracted. These gene tree distributions reflect the topology distributions that were passed to TASTI.
